# CImP: Cellular Imprinting Proteomics applied to ocular disorders elicited by Congenital Zika virus Syndrome

**DOI:** 10.1101/648600

**Authors:** Livia Rosa-Fernandes, Raquel Hora Barbosa, Maria Luiza B. dos Santos, Claudia B. Angeli, Thiago P. Silva, Rossana C. N. Melo, Gilberto Santos de Oliveira, Bernardo Lemos, Martin R. Larsen, Claudete Araujo Cardoso, Giuseppe Palmisano

## Abstract

**IMPORTANCE:** Ocular complications in infants with Congenital Zika Syndrome (CZS) have been reported. However, the molecular mechanisms underlying of eye dysfunctions are presently unknown.

**OBJECTIVE:** A method (termed Cellular Imprinting Proteomics, CImP) for the identification and quantification of the ocular surface proteome using a minimally invasive membrane filter device is described. Moreover, The CImP method was applied to profile the molecular alterations in the eyes of infants exposed to Zika virus (ZIKV) infection during gestation.

**DESIGN, SETTINGS AND PARTICIMPANTS:** The CImP method was applied to a cohort divided into three conditions: 1) Ctrl (infants with no infectious diseases, n=5). 2) Zikv (infants exposed to ZIKV gestation, with no microcephaly, n=5). 3) Zikv^CZS^ (infants exposed to ZIKV, with microcephaly, n=3). All conditions were age and sex-matched. An improved impression cytology method was used to capture the outermost ocular surface cells. The number of impression cytology membrane collected was: Ctrl (12), Zikv (14) and Zikv^CZS^ (8). Proteins were extracted and analysed using mass spectrometry-based proteomics technology followed by statistical analysis. Parallel reaction monitoring was performed to validate the expression of specific protein markers.

**RESULTS:** Using the CImP method, 2209 proteins were identified on the membrane-captured conjunctiva epithelial cells. Modulation of neutrophil degranulation, cell death, ocular and neurodevelopment pathways are reported in infants with CZS compared to matched controls. Moreover, the molecular pattern of ocular surface cells retrieved from infants infected during the gestation but with no CZS was different from matched controls.

**CONCLUSIONS AND PERSPECTIVES:** Molecular alterations in the ocular cell surface associated to ZIKV infection with and without CZS complications are reported for the first time. We predict that this method will be introduced successfully in the study of several neurological diseases with the aim to identify novel diagnostic and therapeutic biomarkers.

## Introduction

Zika virus (ZIKV) is a positive-strand RNA virus belonging to the *Flaviviridae* family that is transmitted to humans predominantly by the female *Aedes aegypti* and *Aedes albopictus* mosquitos (1). Moreover, sexual contact, blood transfusions, laboratory and healthcare settings and congenital transmission are also responsible for ZIKV infection (2). The first report of ZIKV infection was in a nonhuman primate in the in 1947 (3). The first ZIKV transmission to humans was reported in Nigeria (4) and subsequently, outbreaks in Micronesia and French Polynesia occurred (5). In 2015, clinicians in the Northeast Brazil correlated ZIKV infection with reports of newborns with microcephaly, congenital malformations, and neurological syndromes (6). Since then, ZIKV infection raised global public health concern due to its transmission potential and the different fetal and neonatal abnormalities designated Congenital Zika syndrome (CZS) (6). The syndrome characteristics include microcephaly, intracranial calcifications, haemorrhage, ocular pathology, spinal cord and peripheral nerve lesions (7, 8). Indeed, besides central nervous system abnormalities, ophthalmic complications have been associated to ZIKV infection in rhesus monkey and infants with CZS (9). Initial studies in mice lacking the type I IFN receptor showed that ZIKV-inoculated mice developed conjunctivitis and panuveitis. Moreover, ZIKV RNA was detected in all eye regions with higher levels in the retinal pigment/choroid complex and in the optic nerve (10). However, no ocular pathologies were observed in congenitally infected mice. Female rhesus monkeys infected with ZIKV early in pregnancy (weeks 6-7 of gestation) recapitulate the CZS observed in humans and ocular dysfunctions have been described (9, 11, 12). Indeed, Zika virus RNA was found in fetal retina, choroid and the optic nerve (12). Infection of rhesus monkeys with the Brazilian strain of Zika virus was associated to periorbital myositis and myodegeneration (9). Several case reports and studies have described ocular pathologies in neonates born to mother infected with ZIKV during pregnancy (13–19). A common pattern of vision-threatening findings has been associated to congenital ZIKV infection and include chorioretinal atrophy, focal pigmentary changes in the macular region and optic nerve abnormalities (20). Moreover, ZIKV antigens were found in the neural retina, iris, choroid and the optic nerve in infants with CZS (21, 22). The association of eye abnormalities with congenital ZIKV infection has been reported not only in infants with microcephaly but also in infants without CZS signs (23–25). Understanding the molecular mechanisms underlining the ocular abnormalities in ZIKV-infected infants will help in better diagnostic and therapeutic solutions. Impression cytology (IC) is a minimally invasive, easy to perform, robust and reliable method that has allowed the detection of abnormalities and monitoring treatment efficacy in ocular surface cells in several eye diseases (26–45). Likewise, IC has been used to evaluate ocular surface abnormalities in systemic diseases such as diabetes, thyroid orbitopathy, cystic fibrosis, celiac disease and seborrheic dermatitis (41–45). Moreover, the conjunctival epithelial cells were sampled from the eyes of several animal models such as rhesus monkeys, dogs, rabbit and pigs (46–48).

This method is based on the application of a filter material to the exposed interpalpebral surface to remove the outermost layer of conjunctival cells. These cells can be subjected to histological, immunochemical and molecular analyses to identify specific cell morphologies, types and deregulated signalling pathways. Different materials and filter devices have been tested, ranging from polycarbonate, nitrocellulose and cellulose acetate to commercial devices such as the Eyeprim™(28, 49, 50), but no major differences in RNA isolation yields were observed between them(51). Recently, the IC method has been coupled to 2D-DIGE to study protein expression in patients with meibomian gland dysfunction and dry eye disease (52). Barbosa *et.al.* has described an optimized impression cytology method using nitrocellulose membrane and eliminating the use of invasive apparatus, which makes it especially suitable for collections of samples from babies and infants (53).

Here we report: i) the development of the cellular imprinting proteomics method (CImP) for the analysis of the proteome of ocular surface cells and fluid imprinted on a membrane, ii) the application of the CImP method to ZIKV-infected infants with and without CZS and iii) the possibility to identify proteins that could discriminate ocular dysfunctions in infants exposed to ZIKV during gestation.

## Subjects, materials and methods

### 1. Patient cohort

This study comprises 13 infants with and without CZS referred to the Pediatric Service of the Antonio Pedro University Hospital, Fluminense Federal University, Niteroi, Brazil. This study was approved by Institutional review board and ethics committee of Fluminense Federal University, protocol CAAE number 79890517.6.0000.5243, and followed the guidelines of the Declaration of Helsinki. All samples were collected upon informed and written consent from the parents/legal guardians of each participant. Clinical examination was performed by a multidisciplinary team and all infants included in this study are part of a clinical follow up program currently in progress (54). CZS clinical diagnosis was based on the Brazilian Ministry of Health guidelines (55). Retrieved information has been assembled in **Tables 1 and 2**.

**Table 1:** Clinical and demographic data of the patient cohort. Ctrl: children with normal development, without ZIKV exposure. Zikv: children infected by Zika virus during gestation and without Congenital Zika Syndrome. Zikv^CZS^: children infected by Zika virus during gestation and with Congenital Zika Syndrome.

|  | Ctrl | Zikv | Zikv<sup>CZS</sup> |
| --- | --- | --- | --- |
| N° of patients | 5 | 5 | 3 |
| Age at sample collection (months, Mean ± SD) | 21,4 ± 1,8 | 22,4 ± 3,5 | 22,3 ± 4,2 |
| Gender |  |  |  |
| M | 3 | 3 | 2 |
| F | 2 | 2 | 1 |
| Sampling (n° of membranes/subject) |  |  |  |
| 1 | 0 | 1 | 0 |
| 2 | 4 | 2 | 2 |
| 3 | 0 | 1 | 0 |
| 4 | 1 | 0 | 1 |
| 6 | 0 | 1 | 0 |
| Total | 12 | 14 | 8 |
| ZIKV exposition |  |  |  |
| Y | 0 | 5 | 3 |
| N | 5 | 0 | 0 |
| ZIKV symptoms (trimester) |  |  |  |
| 1° | - | 1 | 2 |
| 2° | - | 1 | 1 |
| 3° | - | 2 | 0 |
| Zika congenital infection |  |  |  |
| Y | 0 | 0 | 3 |
| N | 5 | 5 | 0 |
| Microcephaly |  |  |  |
| Y | 0 | 0 | 3 |
| N | 5 | 5 | 0 |
| Vision impairment |  |  |  |
| Y | 0 | 0 | 3 |
| N | 5 | 5 | 0 |
| Type | - | - | optic disc excavation and pallor;<br>abnormal pigment deposition<br>and macular atrophy |

**Table 2:** Clinical and demographic data of the patient cohort and respective mothers. Ctrl: children with normal development, without ZIKV exposure. Zikv: children infected by Zika virus during gestation and without Congenital Zika Syndrome. Zikv^CZS^: children infected by Zika virus during gestation and with Congenital Zika Syndrome.

|  | Ctrl | Zikv | Zikv<sup>CZS</sup> |
| --- | --- | --- | --- |
| Birth date (semester) |  |  |  |
| 1°/2016 | 2 | 4 | 1 |
| 2°/2016 | 3 | 1 | 2 |
| Encephalic perimeter<br>(cm, Mean ± SD) | 33,4 ± 2,6 | 33,6 ± 1,5 | 29,5 ± 0,8 |
| Apgar score |  |  |  |
| 0-3 | 0 | 0 | 0 |
| 4-6 | 0 | 1 | 0 |
| 7-10 | 5 | 4 | 3 |
| Age mother<br>(years, Mean ± SD) | 29 ± 9,1 | 29 ± 6,5 | 33,6 ± 12 |
| Exanthema |  |  |  |
| Y | 5 | 5 | 3 |
| N | 0 | 0 | 0 |
| PCR Zika mother |  |  |  |
| Positive | 0 | 5 | 3 |
| Negative | 5 | 0 | 0 |

### 2. Cellular imprinting proteomics method design

#### 2.1. Ocular cells collection and protein extraction on nitrocellulose membrane

Ocular surface cells were obtained through optimized impression cytology procedure of both eyes as previously described (53). The cell capture area of the nitrocellulose membrane was immersed in a Low-binding Eppendorf tube containing 200µl of protein extraction solution consisting of 1% sodium deoxycholate (SDC), 1× PBS and 1× protease inhibitor cocktail (Sigma Aldrich) and incubated for 20 min under agitation at room temperature. Subsequently, the nitrocellulose membrane immersed in the extraction solution was probe tip sonicated using for three cycles for 20s and intervals of 10s on ice. Left and right eyes were sampled and each membrane sample was analysed separately (**Table 1**).

#### 2.2. Protein reduction, alkylation and trypsin digestion

Proteins were reduced with 10mM DTT for 30min at 56°C and alkylated with 40mM IAA for 30 minutes at room temperature, in the dark. Proteins were quantified using nanodrop method before porcine trypsin (Promega) was added to 1:50 ratio. The digestion, which proceeded for 16 hours at 37°C, was blocked by adding 1% TFA final concentration before the SDC was removed from the solution by centrifugation at 10000xg for 10min. Tryptic peptides were desalted using R3 microcolumns before LC-MS/MS analysis (56, 57).

#### 2.3. Liquid chromatography mass spectrometry analysis

The peptide samples were loaded on an in-house packed pre-column (4cm × 100μm inner diameter, 5μm particles) using an Easy-nanoLC system (ThermoFisher) and separated by gradient from 3 to 28% solvent B in 100 min, 28 to 45% in 20 min, 45 - 95% B in 2 min and 8 min at 95% B. (A = 0.1% FA; B = 90% ACN, 0.1% FA) at a flow of 250nL/min on analytical Reprosil-Pur C18-AQ column (20cm × 75μm inner diameter, 3μm particles).

The Easy-nanoLC system was connected online to Orbitrap Fusion LumosTribrid mass spectrometer (Thermo Fisher) operating in positive ion mode and using data-dependent acquisition. The full MS scans were acquired over a mass range of m/z 350–1600 with detection in the Orbitrap at 120000 resolution with AGC target set to 3e6 and a maximum fill time of 100ms. Following each MS scan, the 20 most abundant peptide ions above a threshold of 50000 were selected in the quadrupole with an isolation window of 0.7 m/z and fragmented using HCD fragmentation (collision energy: 35). Fragment ions were acquired in the orbitrap at 30000 FWHM resolution for an ion target of 50000 and a maximum injection time of 50ms, dynamic exclusion of 30 s and fixed first mass 110 m/z. All data were acquired with Xcalibur software v3.0.63 (Tune v2.0 1258).

#### 2.4. Database search and statistical analysis

Raw data were searched using the MaxQuant v1.6.2.10 (MQ) and Proteome Discoverer v2.3.0.523 (PD) computational platforms using Andromeda (MQ) and Sequest search engines, respectively. The parameters used for database search were: human reviewed proteome database (20400 entries, downloaded from Uniprot the 01/2019) with the common contaminants, trypsin as cleavage enzyme, two missed cleavages allowed, carbamidomethylation of cysteine as fixed modification, oxidation of methionine and protein N-terminal acetylation as variable modifications. Protein identification was accepted with less than 1% FDR. For the Proteome Discoverer platform, the percolator, peptide and protein validator nodes were used to calculate PSMs, peptides and proteins FDR, respectively. FDR less than 1% was accepted. Protein grouping was performed using the strict parsimony principle. Label-free quantification was performed in the two platforms using the extracted ion chromatogram area of the precursor ions activating the matching between run feature. Protein quantification normalization and roll-up was performed using unique and razor peptides and excluding modified peptides. The Intensity based absolute quantification feature (iBAQ) was activated in MaxQuant to calculate the relative protein abundance within samples. Differentially regulated proteins between the three conditions were selected using t-test with a post-hoc background-based adjusted p-value<0.05 (10.1021/pr4006958). Statistical analyses, volcano and PCA plots were performed in the Perseus and Proteome Discoverer software. The data obtained from Proteome Discoverer were used as primary data and complemented with the MaxQuant data to prioritize proteins and biological processes.

#### 2.5. Bioinformatic analysis

The total identified proteins were compared with the human plasma proteome database. Regulated proteins were matched against the human protein atlas database using eye-enriched and brain-enriched proteins (https://www.proteinatlas.org/). Moreover, the NEIBank was used to retrieve disease genes that affect vision and other disease genes with eye phenotypes (https://neibank.nei.nih.gov/cgi-bin/eyeDiseaseGenes.cgi). A total of 441 and 165 genes associated to eye diseases and other diseases were retrieved, respectively. The human plasma proteome was retrieved from the Human Plasma Peptide Atlas, which contains 3509 proteins (http://www.peptideatlas.org/hupo/hppp/). Gene ontologies categories were retrieved using the Protein annotation node built-in Proteome Discoverer. Moreover, STRING database (https://string-db.org/) was used to build protein-protein interaction networks from regulated proteins and identify enriched Reactome and KEGG pathways at FDR<0.05.

### 3. Protein abundance validation using parallel reaction monitoring (PRM)

Differentially regulated proteins were selected for further validation by parallel reaction monitoring using the following criteria: a) protein differentially regulated with adjusted p-value <0.05, b) with more than or equal to 2 peptides, c) with more than or equal to 1 unique peptide, d) concordant abundance ratio of the peptides without missing cleavage and e) more than or equal to 2 unique peptides without missing cleavages found in more than 50% samples of each condition. Parallel reaction monitoring was performed using the same nLC-MS/MS setup described for the discovery experiment including a targeted mass list of peptides selected based on the characteristics described above. A list of peptides, m/z and z is provided (**Supplementary Table S1**). The raw data were searched using Proteome Discoverer (Thermo) as described above for the samples acquired in data dependent mode. The peptide abundances were retrieved from the extracted ion chromatogram (XIC) and normalized by the total base peak intensity for each sample. T-test statistic was performed to calculate differentially regulated peptides (p-value<0.05) between the three conditions. ROC curves and the associated statistical parameters (cut-off, sensitivity and specificity) were calculated using Metaboanalyst computational platform (https://www.metaboanalyst.ca) within the biomarker analysis pipeline. The multivariate ROC analysis was performed using Random Forest as classification and ranking method.

### 4. Morphological analysis and image acquisition

To evaluate microscopic features of the cells, the nitrocellulose membranes containing collected cells were fixed in 4% paraformaldehyde in phosphate buffer, pH 7.4 for 4 h, at room temperature; rapidly vortexed to allow release of adhered cells and cytocentrifuged (Cytospin 4 Shandon, Thermo Scientific, Waltham, MA) at 1200 rpm, for 10 minutes at room temperature. Slides were prepared in quadruplicate from 3 patients per group. For each pair of slides, one was stained with a Diff-Quik kit, as the standard procedure, and the other one with 0.5% toluidine blue O solution (Fisher Scientific) for 5 min. Slides were analyzed on a BX-60 Olympus microscope equipped with an Olympus XC50 CCD camera with cellSens® standard digital imaging software (Olympus Imaging Corp., Tokyo, Japan). A total of 423 cells were analyzed (n= 103 for control group; n= 150 for ZIKV; n= 170 for ZIKV/CZV) for qualitative and quantitative evaluation of morphological alterations. Keratinization was scored as initial to mild (cells showing initial signs of acidophilia at their surface or acidophilia detected in the peripheral cytoplasm) or moderate to severe (cells with a deep cytoplasm partially or fully acidophilic).

## Results

### Clinical features of the patient cohort

Here, the cohort was divided in three conditions: 1) Ctrl, infants with no infectious diseases included in the screen nor neurological disorders without maternal ZIKV exposure; 2) Zikv, infants exposed to ZIKV with no microcephaly; 3) Zikv^CZS^, infants exposed to ZIKV with microcephaly and clinical CZS symptoms. In both Zikv and Zikv^CZS^ conditions, the mothers were ZIKV positive during the first or second trimester of gestation and negative to other infectious agents (syphilis, toxoplasmosis, rubella, cytomegalovirus and HIV) tested. All groups were age and sex matched. The number of subjects was 5, 5 and 3 for the Ctrl, Zikv and Zikv^CZS^ conditions, respectively. Vision impairment was observed only for the three infants belonging to the Zikv^CZS^ conditions at the time of sample collection. The main pathologies observed were optic disc excavation and pallor, macular atrophy and abnormal ocular pigment deposition. Two infants were affected bilaterally and one was affected unilaterally. The ocular surface sampling was performed for both eyes varying between 1 to 6 membranes for each subject. The total number of samples was 12, 14 and 8 for the Ctrl, Zikv and Zikv^CZS^, respectively (**Table 1**). A total of 34 samples were analysed by the CImP method.

### Cellular imprinting proteomics (CImP) method development

The CImP method combines the impression cytology sample collection and a streamlined optimized sample preparation with a large-scale label free quantitative bottom-up proteomics approach (**Figure 1**). The membrane material containing ocular surface cells were transferred to a PBS-buffered solution containing sodium deoxycholate (SDC) detergent and proteases inhibitors. Cells were lysed using probe tip sonication and the proteins digested using trypsin. After tryptic digestion, SDC was removed by acid precipitation and peptides were analysed by LC-MS/MS. The analysis revealed 2209 protein groups identified with at least 1 unique peptide from a total of 17229 peptides (**Supplementary Table S2**). Of them, 2062 proteins were identified in all three conditions (**Supplementary Figure S1A**). Proteins quantified were 865, 1465 and 1842 in 100%, 80% and 50% of the total samples, respectively (**Supplementary Figure S1B**). Peptide abundances were used to derive the principal component analysis. Discrete separation between the three conditions was achieved with the Zikv^CZS^ separated from the others (**Supplementary Figures S1C and S1D**). These proteins were associated to diverse cellular components and biological functions (**Supplementary Figures S2A and S2B**). Response to stimulus and transport were overrepresented in these data. Proteins associated to membrane and extracellular space compartments were also overrepresented, showing a diverse distribution of biological processes and cellular compartment in the conjunctiva epithelial cell proteome. The Intensity Based Absolute Quantification (iBAQ) feature allowed us to quantify the protein abundance of the conjunctiva epithelium within each sample (58). Six proteins, lipocalin-1, lacritin, serum albumin, proline-rich protein 4, lysozyme and mammaglobin-B constitute 50% of the total abundance of the conjunctiva epithelium retrieved by the impression cytology method (**Supplementary Figure S3A**).

**Figure 1:**
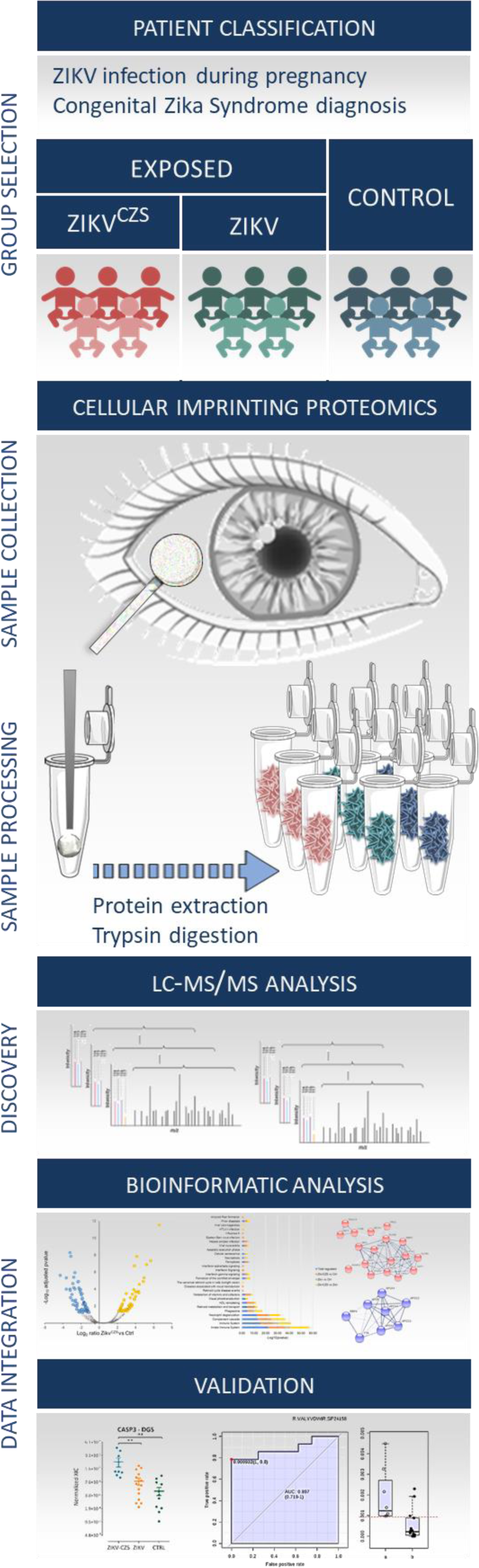
Cellular imprinting proteomics (CImP) method applied to ocular surface alterations during Congenital Zika Syndrome. Experimental workflow of the CImP method applied to a cohort of infants exposed to ZIKV during gestation with Congenital Zika Syndrome (Zikv^CZS^ n=3) and without Congenital Zika Syndrome (Zikv, n=5) and age and sex-matched controls (Ctrl, n=5) (group selection). The sample collection was obtained using nitrocellulose filters used to capture the ocular surface epithelial cells and fluid (sample collection). Proteins were extracted from the membrane, digested and analysed by mass spectrometry (sample processing). Database search and statistical analysis were performed in order to identify differentially expressed proteins (discovery). Protein candidates were validated by parallel reaction monitoring and microscopy analysis confirmed the presence of specific biological processes (data integration).

In order to identify the origin of the conjunctival epithelium proteome retrieved by impression cytology, we compared it with proteomes from different ocular fluids and tissues including tears, aqueous humour, lens, vitreous humour, retina, and retinal pigment epithelium (RPE)/choroid (59). The highest overlap was observed with the RPE/choroid complex proteome. However, vitreous humour, cornea, retina and iris proteomes covered more than 70% of the conjunctiva epithelial proteome. Lens and aqueous humour proteomes had lowest overlap with the conjunctiva epithelium proteome (**Supplementary Figure S3B**). These data indicate that the CImP method can be used in combination with other ocular fluids to discover biomarkers of ocular diseases. In the conjunctival epithelial proteome, we detected blood proteins such as albumin, immunoglobulin, haemoglobin, apolipoprotein, serotransferrin, complement factors and alpha-1-antitrypsin (**Supplementary Table S2**). However, their abundance did not hinder the identification of kinases, receptors and transcription factors involved in several biological processes ranging from metabolism to intracellular signalling (**Supplementary Table S2**). Due to that, we compared the conjunctiva surface proteome with the plasma proteome. In order to identify the human conjunctiva epithelial cells proteins detected in the human plasma, we compared our protein list with the Human Plasma Peptide Atlas containing 3509 proteins (60). There were 1579 proteins common to both conjunctiva epithelial cells and plasma and 630 proteins unique to the ocular cells (**Supplementary Figure S3C**). These data suggest the possibility of using the CImP method to find biomarkers both for ocular and systemic diseases.

### CImP allows the identification and quantification of ocular surface proteins in ZIKV-infected infants

Comparison between the ocular surface proteome versus the brain-enriched and eye-extended proteome retrieved from the human protein atlas, revealed an overlap of 29 and 3 proteins, respectively (**Supplementary Figure S3D**). A comparison of the conjunctiva epithelial proteome with the NEIBank for eye disease genes revealed 66 genes associated to ocular diseases and 23 to other diseases. These data suggest the potential of the CImP method to extract proteins that could mirror ocular and neurological defects (**Supplementary Figures S3E and S3F**). Due to that, we applied the CImP method to identify proteins differentially regulated between the Zikv^CZS^, Zikv and Ctrl conditions. Three comparisons were made between the three conditions: a) Zikv^CZS^ vs Ctrl, b) Zikv vs Ctrl and c) Zikv^CZS^ vs Zikv. The analysis revealed 182 proteins differentially regulated between the Zikv^CZS^ and control with adjusted p-value≤0.05 (**Supplementary Table S3**). In total 82 and 100 proteins were up and downregulated, respectively (**Figure 2A, Supplementary Table S3**). A total of 166 proteins were found differentially regulated between the Zikv and control with adjusted p-value≤ 0.05. In total 70 and 96 proteins were up and downregulated, respectively (**Figure 2B, Supplementary Table S4**). Lastly, 204 proteins were differentially regulated between the Zikv^CZS^ and Zikv with adjusted p-value≤ 0.05. In total 110 and 94 proteins were up and downregulated, respectively (**Figure 2C, Supplementary Table S5**). Combining the differentially regulated proteins in the three comparisons, 340 proteins were identified. Protein-protein interaction networks analysis of the different comparisons revealed 70 and 14 Reactome pathways enriched in the Zikv^CZS^ vs Ctrl and Zikv vs Ctrl comparisons, respectively (**Supplementary Figure S4A**). The most enriched processes were associated to a) immune system, b) ocular dysfunctions, c) interferon signalling, d) cell death, e) viral infection and f) neurological disorders (**Figure 3**). The immune system, neutrophil degranulation, visual phototransduction, metabolism of retinoids, interferon signalling, viral infection processes were mostly enriched in the Zikv^CZS^ vs Ctrl and Zikv^CZS^ vs Zikv comparisons. Selected proteins involved in these processes were validated by PRM analysis.

**Figure 2:**
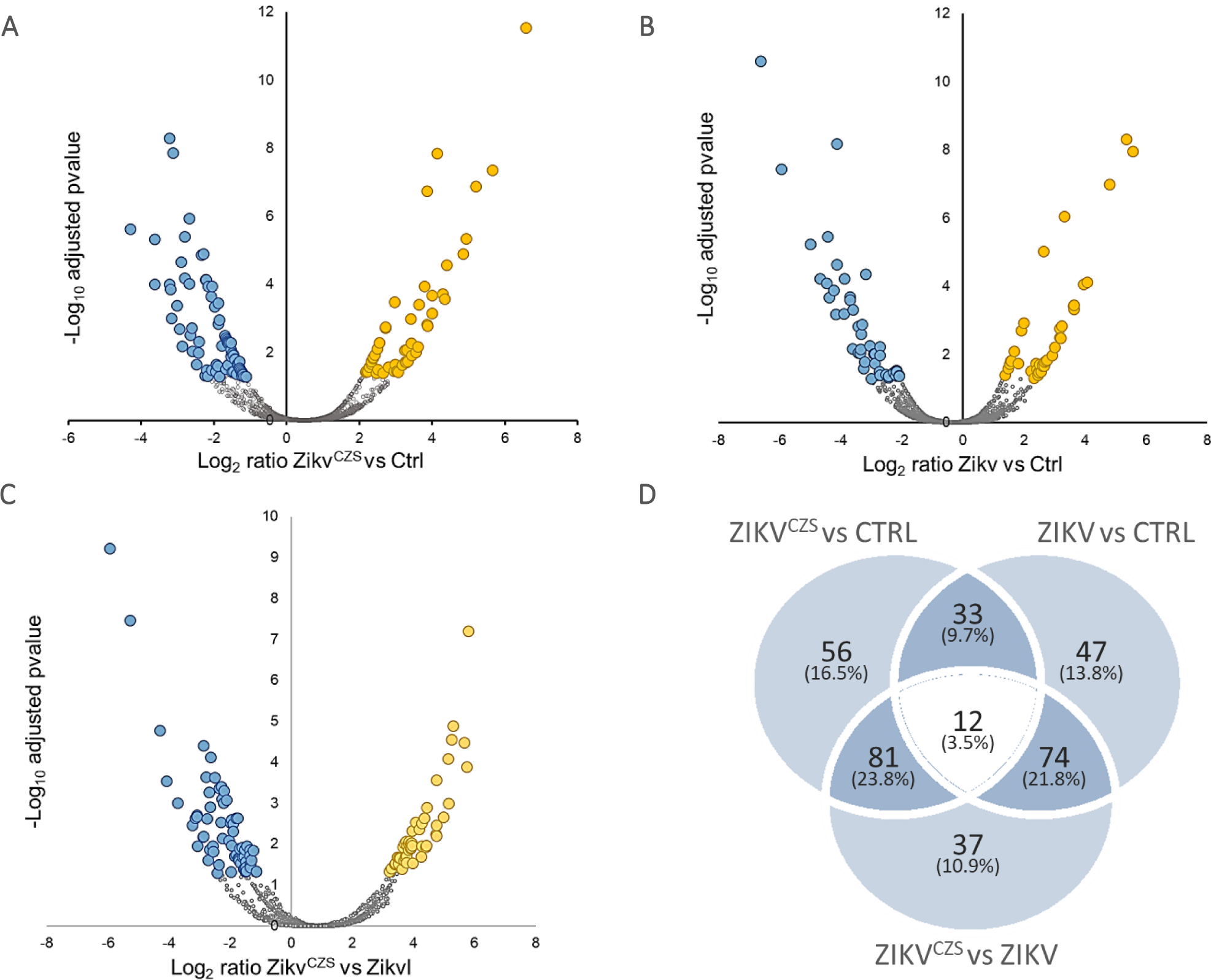
Proteins differentially regulated between Zikv^CZS^ vs Ctrl (A), Zikv vs Ctrl (B) and ZikvCZS vs Zikv (C). Proteins with adjusted pvalue<0.05 were considered statistically significant. D) Overlap between the differentially regulated proteins in the three comparisons.

**Figure 3:**
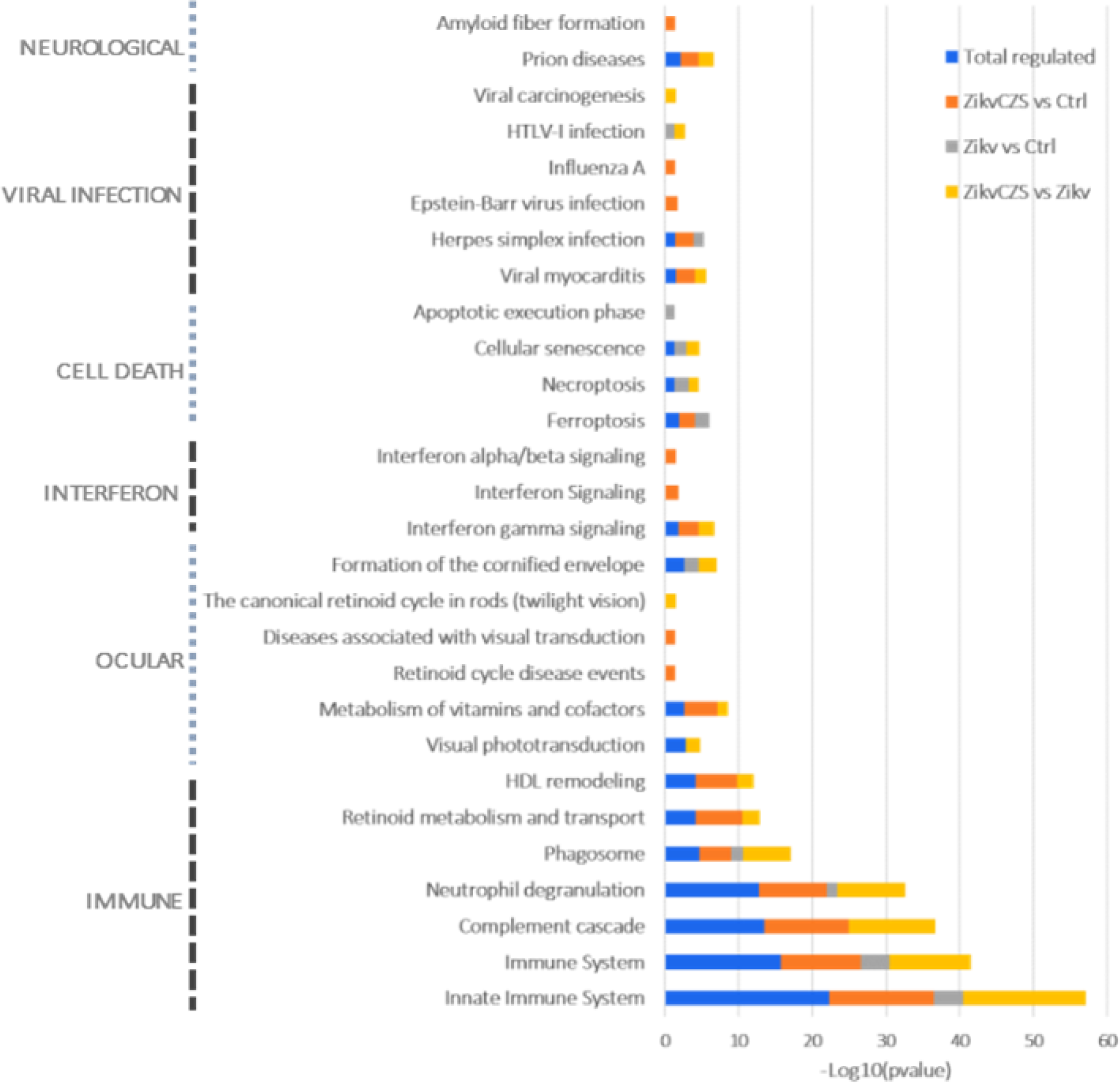
Pathways modulated in Zikv^CZS^, Zikv and Ctrl conditions. Pathways enriched in the different datasets (total regulated, Zikv^CZS^ vs Ctrl, Zikv vs Ctrl and Zikv^CZS^ vs Zikv) and grouped into immune system, ocular disorders, interferon signalling, cell death, viral infection and neurological diseases. B) Differentially regulated proteins associated to the ocular disorders pathway identified during the discovery phase and validated by PRM. The peptide abundance for each protein is reported for the Zikv^CZS^, Zikv and Ctrl groups.

### Validation of differentially regulated proteins involved in immune response, ocular disorders, cell death and neurological diseases

Selected proteins associated to immune system response, ocular disorders and cell death and identified as differentially regulated during the discovery phase were validated by parallel reaction monitoring. Two peptides per protein were monitored. A total of 72 precursor ions, corresponding to 54 peptides and 26 proteins were monitored (**Supplementary Table S1**). Proteins were considered validated when two peptides belonging to the same protein were statistically significant between the conditions and with the same regulation trend, except for gasdermin-D, MMP8 and S100A12, proteins that were detected with one unique peptide (**Supplementary Table S6**). The comparison between Zikv^CZS^ and Ctrl confirmed the regulation of MPO, ELANE, ITGAM, RNASE3, AZU1, MMP8, PRTN3, ALDH3A1, CASP3, BID, GSDMD, DEFA1, S100A12, CPNE1, EPS8L2 and DMBT1 (**Supplementary Table S6**). The comparison between Zikv and Ctrl confirmed the regulation of HIST1D, BASP1 and DMBT1 (**Supplementary Table S6**). The comparison between Zikv^CZS^ and Zikv conditions confirmed the regulation of MPO, ELANE, ITGAM, AZU1, PRTN3, DEFA1, S100A12, CPNE1 and DMBT1 (**Supplementary Table S6**). The single peptide abundances were used to calculate the ROC curve, sensitivity and specificity to discriminate the three conditions. AUC more than 0.8 on the two peptides belonging to each protein were considered as potential discriminative features. For the Zikv^CZS^ vs Ctrl comparison, peptides belonging to ALDH3A1, CASP3, DMBT1, ITGAM, MMP8, MPO, PRTN3 and RNASE3 proteins fulfilled the criteria (**Supplementary Table S6, Supplementary Figure S5**). All proteins were upregulated in the Zikv^CZS^ condition compared to control. For the Zikv vs Ctrl comparison, peptides belonging to HIST1H1D and BASP1 were downregulated in the Zikv condition compared to control. (**Supplementary Table S6, Supplementary Figure S6**). For the Zikv^CZS^ vs Zikv comparison, peptides belonging to AHNAK, AZU1, CASP3, DEFA1, DMBT1, ELANE, ITGAM, MPO, PRTN3, RETN, RNASE3, S100A12 (**Supplementary Table S6, Supplementary Figure S7**). All these proteins were upregulated in the Zikv^CZS^ condition compared to Zikv. A larger dispersion of the Zikv^CZS^ samples was observed compared to the other conditions indicating a variability of the syndrome in the clinical and molecular readout.

### Neutrophil degranulation signature and immune response in the eye

The most enriched molecular pathway throughout the comparisons was immune system response (**Figure 3**). In particular, 19 proteins were associated to neutrophil degranulation and all of them were upregulated in the Zikv^CZS^ compared to controls, indicating an activation of this process (**Supplementary Figure S4B**). Indeed, a neutrophil protein signature based on neutrophil defensin 1 (DEFA1), neutrophil collagenase (MMP8), cytochrome b-245 heavy chain (CYBB), resistin (RETN), DnaJ homolog subfamily C member 5 (DNAJC5), olfactomedin-4 (OLFM4), eosinophil cationic protein (RNASE3), ADP-ribosyl cyclase/cyclic ADP-ribose hydrolase (BST1), protein S100-A12 (S100A12), neutrophil elastase (ELANE), azurocidin (AZU1), pentraxin-related protein (PTX3), chitotriosidase (CHIT1), Bone marrow proteoglycan (PRG2), gamma-glutamyl hydrolase (GGH), chitinase-3-like protein 1 (CHI3L1), protein S100-A7 (S100A7), neutrophil cytosol factor 2 (NCF2) and myeloperoxidase (MPO) was found in Zikv^CZS^ group (**Figure 4**). The upregulation of MPO, ELANE, ITGAM, RNASE3, AZU1, MMP8, PRTN3, DEFA1 and S100-A12 were validated by parallel reaction monitoring (**Supplementary Table S6, Figure 4**). The interferon signalling pathways were enriched in the regulated proteins belonging to the Zikv^CZS^ vs Ctrl comparison (**Figure 3**). IFIT3 protein was significantly upregulated 38 times in the Zikv^CZS^ group compared to controls (**Supplementary Table S2**).

**Figure 4:**
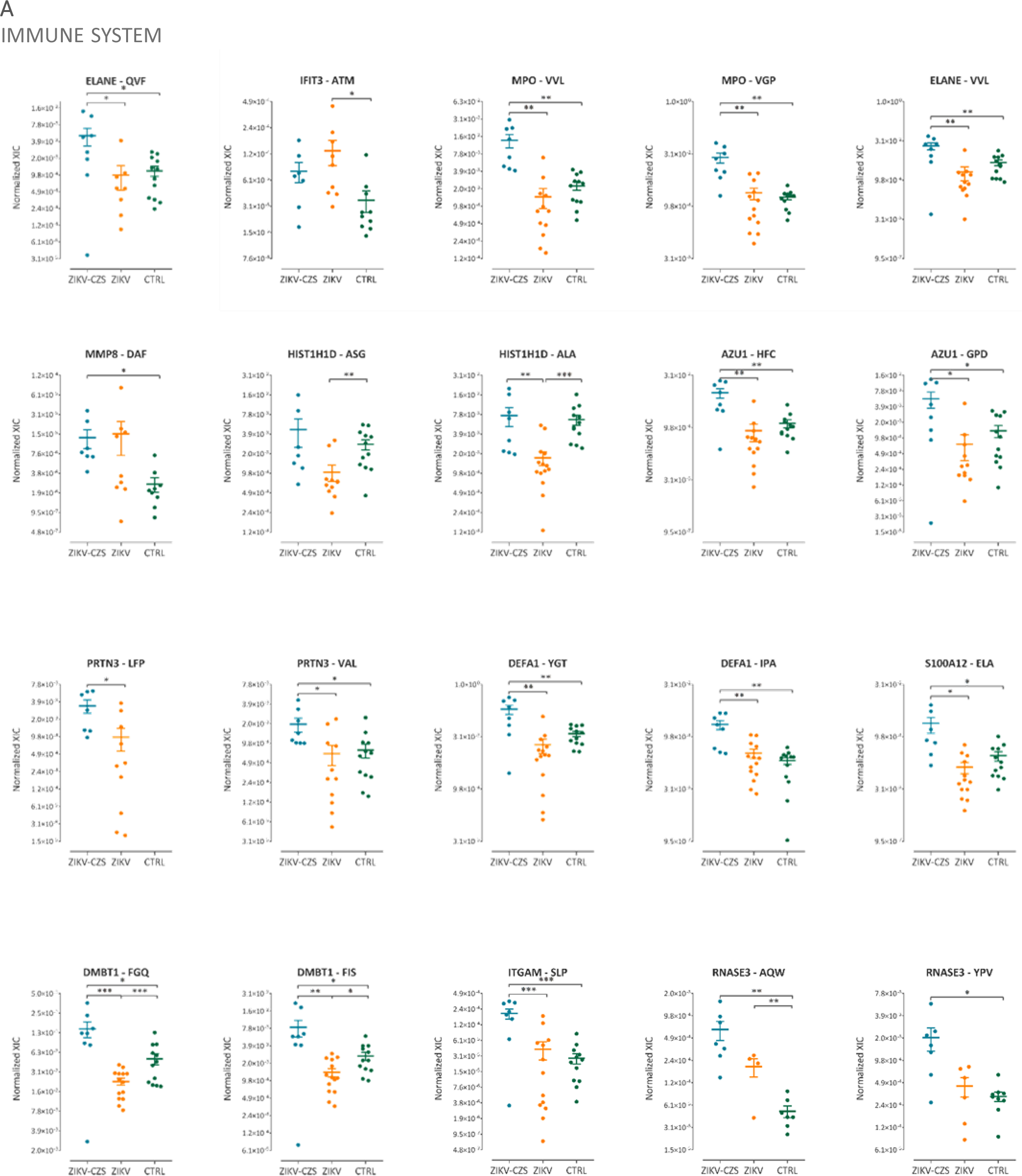
Differentially regulated proteins belonging to the immune system pathway identified during the discovery phase and validated by PRM. The peptide abundance for each protein is reported for the Zikv^CZS^, Zikv and Ctrl groups.

### Activation of cell death in the conjunctiva epithelium of Zikv-infected infants

Cell death pathways were enriched in the regulated proteins between the Zikv^CZS^ and control and the Zikv and control conditions (**Figure 3**). Gasdermin D was identified to be upregulated 6.5 times in the Zikv^CZS^ condition at p-value<0.01 and adjusted p-value<0.07 during the discovery phase. Differential upregulation of Gasdermin D in Zikv^CZS^ compared to control was confirmed by parallel reaction monitoring of the ELCQLLLEGLEGVLR peptide, p-value=0.04 (**Figure 5A**). Moreover, caspase-3 (CASP3) and BH3-interacting domain death agonist (BID) were differentially regulated in the Zikv^CZS^ condition compared to controls being 3.5 and 4 times upregulated in the Zikv^CZS^, respectively (**Figure 5A**). Bright field microscopy of ocular epithelial cells isolated from ZIKV-infected children with CZS show clear morphological evidence (**Figure 5B**) of cell death such as pyknosis (**Figure 5C**), karyorrehesis (**Figure 5D**), karyolysis (**Figure 5E**), cytoplasmic vacuolization (**Figure 5F**) and disintegration of the nucleus (**Figure 5G**). A high percentage of cells with these morphologies were measured in Zikv^CZS^ condition **Figure 5H**).

**Figure 5:**
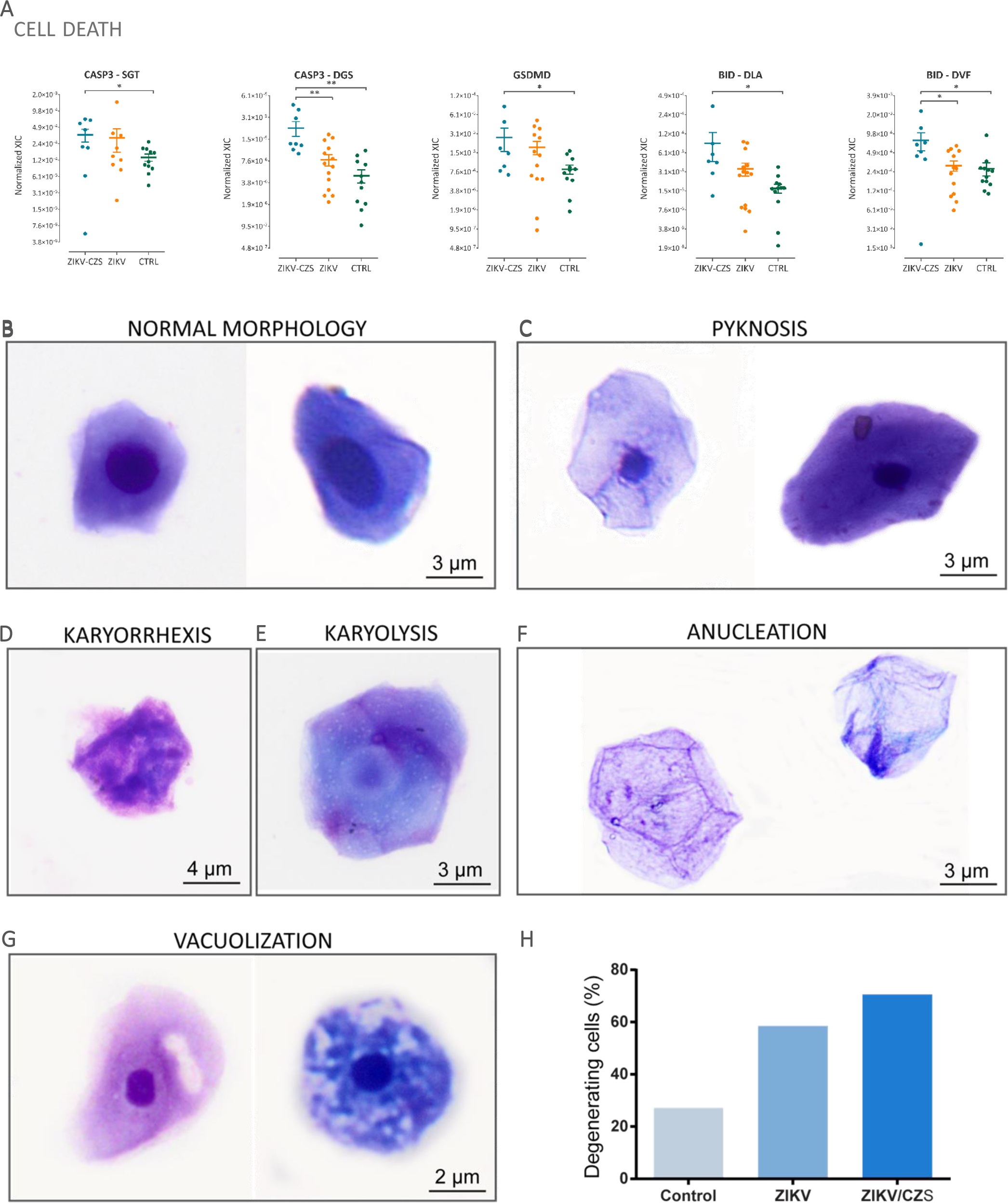
Differentially regulated proteins belonging to the cell death pathway identified during the discovery phase and validated by PRM. The peptide abundance for each protein is reported for the Zikv^CZS^, Zikv and Ctrl groups (A). Representative cells from uninfected group show normal morphology (B). Nuclear and cytoplasmic alterations observed in cells collected from ZIKV (infected with no clinical signs) (B–left panel; C and F, left panel) or ZIKV/CZV (diagnosed congenital Zika syndrome (B, right panel; D, E and F, right panel) children. Quantitative analyses show increased number of conjunctval cells with morphological signs of degeneration/cell death. Cells were collected by impression cytology, cytocentrifuged and stained with Diff-Quik or toluidine blue (H). A total of 423 cells were counted and scored for morphological changes.

### Differential regulation of proteins associated to ocular diseases

Ocular processes were enriched using proteins differentially regulated between the three conditions (**Figure 3 and Supplementary Figure S4C**). In our study, proteins associated with retinoid metabolism were downregulated in Zikv^CZS^ compared to controls (**Supplementary Figure 4C**). Several proteins belonging to this pathway were validated by PRM (**Figure 6A**). Retinol-binding protein 4 (RBP4) was found downregulated by more than 50 times in Zikv^CZS^ compared to control. The YWGVASFLQK peptide belonging to RBP4 was found downregulated in the Zikv^CZS^ compared to control conditions (**Supplementary Table S6**). Furthermore, six apolipoproteins (APOA1, APOA2, APOA4, APOB, APOC2 and APOC3) were downregulated in Zikv^CZS^ compared to controls (**Supplementary Figure S4C**). APOB was downregulated 3 times in the Zikv^CZS^ condition (**Supplementary Table S2**). The upregulation of Aldehyde dehydrogenase dimeric NADP-preferring (ALDH3A1) in the Zikv^CZS^ and Zikv conditions compared to controls could be related to the eye protection against oxidative damage. Alcohol dehydrogenase 1B (ADH1B) was identified downregulated in the Zikv^CZS^ compared to control condition (**Supplementary Table S6**). Proteins associated to the cornified envelope were regulated in the three comparisons. The CImP method allowed the identification of 30 keratin proteins (KRT1, 2, 3, 4, 5, 6A, 6B, 7, 8, 9, 10, 12, 13, 15, 15, 16, 17, 18, 19, 23, 24, 25, 31, 73, 74, 76, 77, 78, 80 and 85, **Supplementary Table S2)**. A different regulation pattern of keratins was identified in the Zikv^CZS^ vs Ctrl (KRT15 and 23), Zikv vs Ctrl (KRT3, 6B, 16, 23, 31 and 76) and Zikv^CZS^ vs Zikv (3, 6B, 16, 23, 31, 76 and 85), (**Supplementary Tables S3, S4 and S5)**. Several keratinization-related proteins were regulated. Reflecting the proteomics findings, quantitative analysis by bright field microscopy of conjunctival epithelial cells obtained by impression cytology from children with CZS show a higher degree of moderate to severe keratinization while initial to moderate keratinization can be observed in the Zikv group, and the control group presented less cells displaying any level of this process (**Figure 6B**).

**Figure 6:**
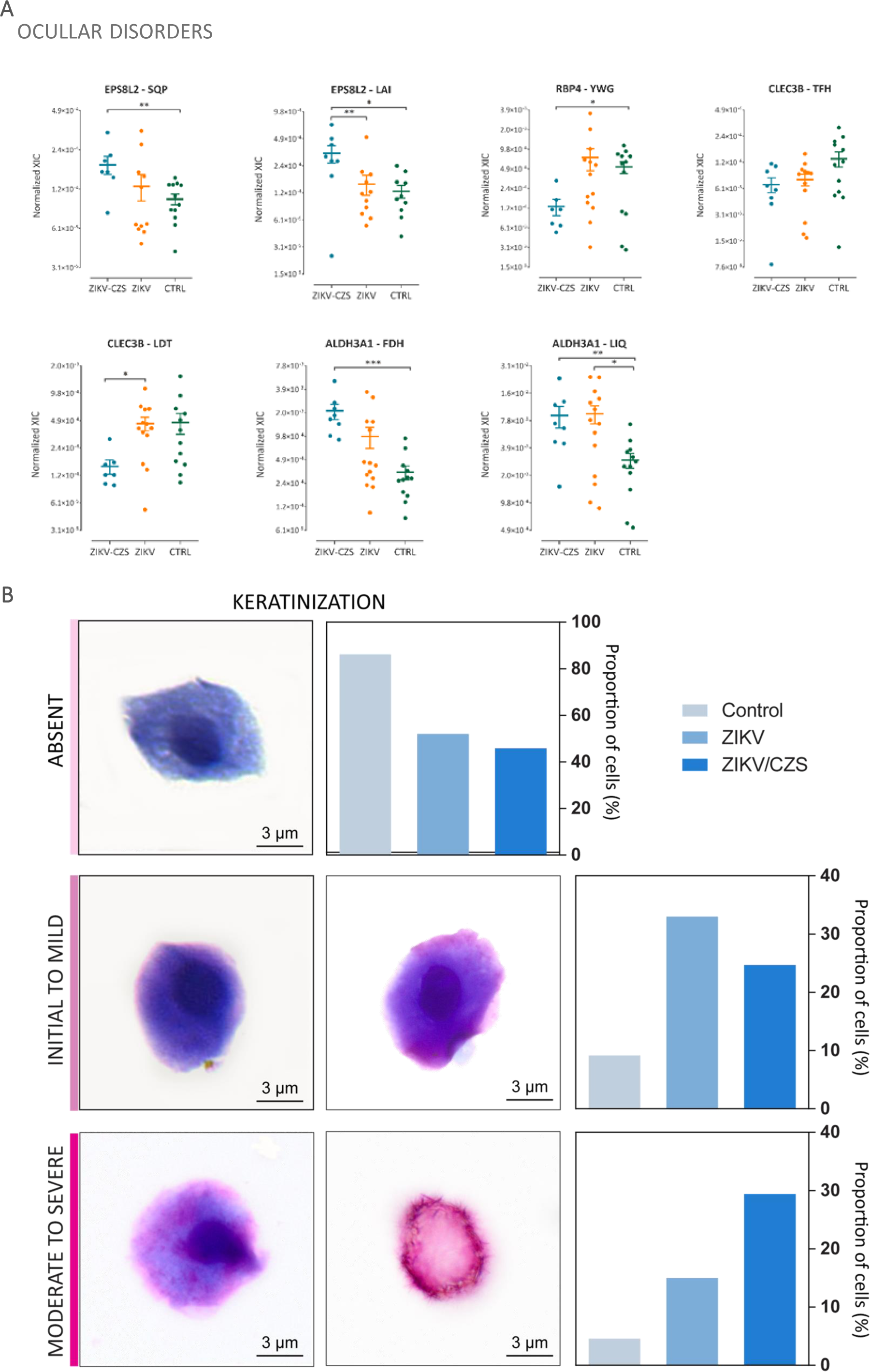
Differentially regulated proteins belonging to the ocular disorders pathway identified during the discovery phase and validated by PRM. The peptide abundance for each protein is reported for the Zikv^CZS^, Zikv and Ctrl groups. Representative images and quantitative morphological analyses of cells collected from children with ZIKV (infected with no clinical signs), ZIKV/CZV (diagnosed congenital Zika syndrome) or uninfected controls using impression cytology. Cytocentrifuged preparations were stained with Diff-Quik (B). A total of 423 cells were counted and scored for morphological signs of Keratinization.

### Neurological diseases-associated proteins

One of the statistically enriched processes was related to neurodevelopment (**Figure 3**). Due to that, we investigated specific proteins differentially regulated in the Zikv^CZS^ vs Zikv comparison that could be related to this process. Copine-1 and Reticulon-4 were upregulated while the Afamin and Pigment epithelium-derived factor were downregulated (**Supplementary Table S2**). Copine-1 was significantly upregulated (p-value<0.001) in the Zikv^CZS^ vs Ctrl upon monitoring by PRM two peptides (SDPLCVLLQDVGGGSWAELGR and EALAQTVLAEVPTQLVSYFR) (**Supplementary Table S6)**. Reticulon-4 (RTN4) was identified upregulated in the Zikv^CZS^ compared to control conditions. Afamin (AFM) was downregulated 4 times in the Zikv^CZS^ group. Pigment epithelium-derived factor (PEDF) was significantly downregulated in the Zikv^CZS^ vs Ctrl comparison.

### Infants exposed to ZIKV infection during gestation and without CZS harbour differential ocular surface proteome

Clinical cases of adults infected by ZIKV presenting ocular abnormalities have been described (61). These data indicate that the neurotropism of the virus is not a necessary factor to elicit ocular dysfunctions. Indeed, in our cohort, the ocular surface proteome of the Zikv group compared to controls showed the regulation of 166 proteins (**Figure 2**). The number of enriched Reactome pathways and their significance had lower p-values was lower compared to the Zikv^CZS^ vs control comparison (**Supplementary Figure S4A**). Immune system response, cell death and cornified envelope proteins were enriched. For the immune response pathway, the IFIT3 protein was upregulated 17 times in the Zikv compared to Ctrl (**Supplementary Table S2**). The upregulation of IFIT3 was confirmed by PRM using the ATMYNLLAYIK peptide in the Zikv group (p-value=0.03), (**Supplementary Table S6)**. Moreover, interferon-induced helicase C domain-containing protein 1 (IFIH1) was upregulated 27 times in the Zikv group compared to Ctrl (**Supplementary Table S2**).

PRM analysis of selected protein candidates identified HIST1H1D and BASP1 as discriminating proteins for the Zikv vs Ctrl conditions with AUC>0.8. These proteins were downregulated in the Zikv condition compared to control (**Supplementary Table S6, Supplementary Figure S6**). None of the children exposed to ZIKV without CZS had visual impairment at the time of collection. However, significant molecular alterations were detected in the eyes of infants exposed to ZIKV.

## Discussion

Here, we present the CImP method, Cellular Imprinting Proteomics, build on the optimized impression cytology procedure previously described (doi: https://doi.org/10.1101/577668) followed by an integrated mass spectrometry-based quantitative proteomics approach to identify the ocular epithelial surface proteome and its modulation in pathophysiological conditions. Impression cytology technique has emerged as an important non-or minimally invasive technique to diagnose and understand molecular alterations in ocular diseases. As of January 2019, 936 manuscripts were retrieved in Medline (PubMedNCBI http://ncbi.nlm.nih.gov) using the term “impression cytology” in the title/abstract and only 5 of them were associated also to the word “proteomics”. One seminal work on impression cytology and proteomics used 2D-DIGE approach to investigate the ocular dysfunctions in patients with meibomian gland dysfunction and dry eye disease was published (52). In this study, an 8 mm in diameter cellulose disc was applied to the eye on both sides to improve the number of captured cells. A lysis buffer containing urea as chaotropic agent and CHAPS as detergent was used to extract cellular proteins before precipitation and resuspension for 2D-DIGE. Using this technique, 348 protein spots corresponding to different proteoforms were visualized. Statistical analysis revealed 17 differentially regulated spots that were selected for MALDI TOF/TOF analysis. Dotblot validation of annexin A1, protein S100A8 and protein S100A4 was performed, showing the reliability of this technique.

Applying the CImP method, protein extraction is based on SDC, a trypsin and MS-compatible detergent, allowed a streamlined approach without extra steps and protein loss. Membranes from the left and right eye of each patient were collected to improve the sampling coverage. The CImP method was applied to a cohort of infants exposed to ZIKV infection during gestation with or without microcephaly grouped into three conditions: Ctrl, Zikv and Zikv^CZS^. Large-scale quantitative proteomics using nLC-MS/MS allowed us to identify 2062 proteins with high confidence, representing the first description of the ocular epithelial proteome collected by impression cytology. Intra-sample quantitative analysis revealed that six proteins (lipocalin-1, lacritin, serum albumin, proline-rich protein 4, lysozyme and mammaglobin-B) constituted 50% of the total amount of the ocular epithelial proteome. Lipocalin, lacritin and lysozyme are also involved in host response to infections. Lipocalin-1 (LCN1, human tears lipocalin) and neutrophil gelatinase-associated lipocalin (LCN2) were among the top 100 most abundant proteins in the ocular surface conjunctiva proteome (**Supplementary Figure S2A**). Lipocalins are extracellular proteins and belong to the lipocalin superfamily that bind and transport hydrophobic compounds such as lipids, hormones and vitamins. LCN1 binds phospholipids and retinol (62) and is expressed mainly in the lachrymal and salivary glands. LCN2 binds preferentially iron and is expressed mainly in the bone marrow (63). Moreover, LCN1 and LCN2 mediate immune and inflammatory response against microbial and fungal infections (64–66). Interestingly, LCN1 was the most abundant protein within the total proteins identified, contributing to 18% of the total protein abundance. Dota A. et al. showed that LCN2 was a conjunctiva epithelium-specific gene (67). The difference could be due to the collection method. Indeed, compared to our improved impression cytology approach, brush cytology was used to obtain conjunctival epithelium, a method that collects all epithelial cell layers of the conjunctival epithelium (67). Human secretoglobin 2A1 (mammaglobin-B, SCGB2A1) is a small protein with unknown function, homolog to mammaglobin-A, that is expressed in tears, lacrimal glands and in the keratinized stratified squamous epithelium of the eyelid as well as in the stratified epithelium of the conjunctiva and in the orbicularis oculi muscle (68). Lysozyme is one of the most abundant proteins in the human tear proteome (1mg/mL) with antimicrobial activity (69). Comparison with the proteome of other ocular tissues and fluids revealed high similarity with the RPE/choroid and less similarity with lens and aqueous humour. Furthermore, the CImP method was applied to map the ocular surface proteome of infants exposed to ZIKV infection with and without CZS compared controls. Proteins differentially regulated between the different comparisons revealed the modulation of pathways involved in immune system response, cell death, ocular and neurological dysfunctions. A total of 26 proteins involved in different cellular events were validated by PRM. Proteins associated to immune system response and neutrophil degranulation were upregulated in the Zikv^CZS^ and Zikv compared to Ctrl condition. The upregulation of neutrophil-associated proteins during flavivirus infection has been shown (70, 71). Neutrophil defensin-1 belongs to the defensins family present in the neutrophil granules with activity against bacteria, fungi, parasites and viruses (72, 73). DEFA1 was found upregulated in the cerebrospinal fluid of patients infected with West Nile virus with neuroinvasive complications (74). DEFA1 gene expression was upregulated in the blood of Dengue-infected children with haemorrhagic fever (75). Myeloperoxidase activity and expression was elevated in mice infected by the intracranial route by DENV-3 confirming the neutrophil infiltration (76). Moreover, MPO was released by neutrophils collected from mice infected with Japanese Encephalitis virus (71). A transcriptomic analysis of peripheral blood mononuclear cells isolated from children with acute-phase Dengue haemorrhagic fever revealed an activation of the S100A12 protein (77). Neutrophil collagenase was released in the culture medium of immune cells treated with Dengue virus NS1 resulting in endothelial glycocalyx degradation and vascular permeability (78). Olfactomedin-4 is a granular protein induced by G-CSF in mature granulocytes (79, 80), also expressed in a subset of neutrophils and was found regulated in septic patients and in paediatric respiratory syncytial virus infection (81). Interestingly, OLFM4 null mice challenged with *Staphylococcus aureus* and *Helicobacter pilori* were more resistant to infection compared to the wild type (82, 83).

As observed for West Nile virus, herpes simplex virus-1 and ebola virus, Zika virus has tropism for immunoprivileged organs like the brain and the eye (84, 85). The infection and persistence in these organs pose new challenges in understanding the dynamics of viral spreading, life cycle and clearance. Neutrophils are essential effectors of the innate immune response. Their recruitment in an infected tissue is a host-defence mechanism that can induce tissue damage if not finely regulated. Infection of neonatal C57BL/6 wild-type mice at 1?day post birth with ZIKV showed that the virus infects the cornea and retina resulting in chorioretinal lesions (86). The infection is followed by inflammatory cells infiltration and increased local expression of chemokines. In ZIKV-infected patients, there is an increased cellular infiltration and inflammation (87). Neutrophil degranulation pathway was found to be activated in the conjunctival epithelial cells retrieved from ZIKV-infected children with CZS. This pathway is associated to neutrophils migration towards the inflammatory site mobilizing granules with antimicrobial properties. Moreover, neutrophils secrete inflammatory cytokines and mediators stimulating T-cells. Immune activation has been associated to inflammatory and degenerative eye diseases such as adult macular degeneration and uveitis (88). In the brain of infected mice, neutrophils and mononuclear cells were infiltrated and polymorphonuclear cells (PMNs) were detected near blood vessel (89). Neutrophil and eosinophil infiltration has been described in several ocular pathologies such as ocular Stevens-Johnson syndrome, mucous membrane pemphigoid, Wegener’s granulomatosis and cicatricial pemphigoid (90–93). Microscopy analysis of the cells content of membranes retrieved from the Zikv and Zikv^CZS^ conditions revealed an increase in immune cells infiltrates associated with an activated ocular surface inflammatory response to ZIKV infection. In particular, we show the overexpression of neutrophils and eosinophil granule constituents correlated to the activation of neutrophil degranulation process in Zikv and Zikv^CZS^ children compared to controls. It is unclear if the immune cells infiltration would be the indirect cause of chorioretinal lesions or if the ZIKV itself elicits these eye abnormalities or a combination of the two phenomena.

Another important cell biology process that interferes with the flavivirus is the interferon signalling pathway. RNA intermediates of viral replication are recognized by cytosolic and endosomal pattern recognition receptors that induce type-I interferon expression and secretion which leads to autocrine and paracrine activation of interferon stimulated genes such as interferon-induced with tetratricopeptide repeats (IFIT) genes. These genes control early viral infection in several cells and tissues. It has been reported that Zika virus upregulates IFIT1 and IFIT3 expression in human dendritic cells (94). Moreover, ZIKV increase the expression of IFIT1 in mouse and human testis (95) and the upregulation of ISG genes has been found in mice brains infected with lymphocytic choriomeningitis virus or West Nile virus (96). These results indicate that ocular surface expression of IFIT3 can be associated to Zika virus infection.

Another important process that was activated in the eyes of infants exposed to ZIKV infection was cell death. Gasdermin is an effector of pyroptosis cell death (97, 98). Pyroptosis was observed in 1992 as Caspase-1-programed cell death and was later termed in 2001 as to differentiate it from the morphologically distinct apoptosis. Age-related macular degeneration involves primarily the cell death of retinal pigment epithelium. Gao J. et al. injected amyloid beta intravitreously inducing sustained activation of the inflammasome (99). They identified the activation of apoptosis and pyroptosis mediated by enhanced caspase-1 immunoreactivity, augmented NF-κB nuclear translocation, increased IL-1β vitreal secretion, and IL-18 protein levels. Moreover, elevated levels of cleaved caspase-3 and gasdermin D were found in RPE-choroid tissues. Moreover, the role of pyroptosis effectors was investigated in in pterygia compared with normal human conjunctival epithelium (100). Pterygium is a common ocular disease characterized by proliferating fibrovascular tissue. Pyroptosis displays a role in the pathological process of pterygium formation and progression. Recently, differential Gasdermin-D expression and cleavage was reported in experimental cerebral ischemia and reperfusion model (101). NLRP3 activation increased the levels of IL-18 and IL-1b after middle cerebral artery occlusion/reperfusion and this process was mediated partly by caspase-1 dependent cleavage of gasdermin-D. The upregulation of GSDMD, CASP3 and BID in the eyes of infants exposed to Zikv during the first trimester of gestation indicate an increase in ocular cell death.

The metabolism of retinoids in the retina is essential for vision (102). Indeed, the photoisomerization of opsin-bound 11-*cis*-retinal to all-*trans*-retinal is critical to activate the phototransduction signalling events within the visual cycle. In order to restore the 11-*cis*-retinal, a series of metabolic reactions are needed including the reduction of all-*trans*-retinal to all-*trans*-retinol (103). All-*trans*-retinol, also known as vitamin A, is also introduced by food intake. Dysregulation of retinoid metabolism has been associated with several ocular diseases such as Stargardt disease, Autosomal recessive cone-rod dystrophy, Autosomal recessive childhood-onset severe retinal dystrophy, Leber congenital amaurosis and Autosomal recessive retinitis punctata albescens (104–107). RBP4 is a 21kDa protein synthesized primarily in the liver and other tissues such as retinal pigment epithelium and choroidal plexus brain (108, 109). It belongs to the lipocalin superfamily and the main function is to bind retinol (Vitamin A_1_) in hepatocytes and deliver it to peripheral tissues (110). Retinol, in the form of retinal, is essential for the visual cycle and defects in RBP4 expression have been associated to ocular diseases. In human patients, iris coloboma, atrophy or focal loss of the retinal pigment epithelium (RPE) and the choroid have been associated to a homozygous splice site variant (c.111+1G>A) and heterozygous missense mutations (Ile41Asn and Gly75Asp) in the gene encoding retinol binding protein 4 and consequent serum level reduction (111, 112). Mice lacking RBP4 had impaired visual function and transgenic expression of human RBP4 induced normal electro-retinogram and retinol metabolism (113–115).

Six apolipoproteins were downregulated in the Zikv^CZS^ condition compared to control. Recently, ApoA-I has been identified as an all-*trans*-retinoic acid-binding protein in the choroid and sclera conditioned media of chick tissue (116). Abetalipoproteinemia is an autosomal recessive disease caused by genetic mutations in the MTTP (microsomal triglyceride transfer protein) gene that result in the inability to synthesize the APOB. The signs and symptoms of this disease are severe deficiency of fat-soluble vitamins such as vitamin A, E and K. The ocular complications of this disease involve diffuse and sometimes patchy pigmentary changes often called atypical retinitis pigmentosa (117). Aldehyde dehydrogenase dimeric NADP-preferring (ALDH3A1) is an enzyme that catalyses the oxidation of aldehydes compromise 5-50% of the human cornea (118–120). ALDH3A1 has a protective function in the cornea by reducing 4-Hydroxynonenal–protein adducts upon exposure to UV radiation (121). Moreover, ALDH3A1 exerts a “suicide” function in the cornea by autoxidation of specific amino acids, protecting it from UV light exposure (121, 122). ALDH3A1 null mice developed cataracts at one-month age (123). One of the ocular abnormalities observed in children infected with Zika virus was cataract formation. Moreover, cornea opacity was detected in ZIKV-infected infants. ALDH3A1 was identified upregulated in pterygium tissues compared to healthy conjunctiva (124). High concentrations of corneal ALDH3A1 have been associated to antioxidant protection of UV-induced free radicals and through the synthesis of NADPH (125, 126). Alcohol dehydrogenase is a zinc metalloenzyme that catalyse the oxidation of alcohol to aldehyde displaying specificity for different substrates such as retinol (127). ADH1B was identified downregulated in keratoconus corneal fibroblasts compared to healthy cornea fibroblasts and suggested as a possible marker of keratoconus (128). Interestingly, ADH1B was not regulated in the comparison between ZIKV and controls.

Several proteins involved in the cornified epithelium envelope were identified. The cornified cell envelope constitutes a protective barrier of squamous epithelial cells (129). In skin keratinocytes, cornified cell envelope precursors such as involucrin, loricrin, small proline-rich proteins, late envelope proteins (LEPs), and filaggrin are crosslinked by transglutaminase forming a 5- to 10-nm mature envelope adjacent to the cell membrane (130–132). Corneal cell envelope bears similarities with the epidermidis (133). However, the absence of a water-impermeable layer and the need of oxygen and nutrients to permeate the cornea epithelium indicate that envelope proteins are associated to other functions besides protective. Pathological keratinization of the corneal and conjunctival mucosal epithelia is associated to ocular diseases and severe visual loss (132). Elevated levels of involucrin and filaggrin have been associated to Stevens-Johnson syndrome (132). Moreover, upregulation of transglutaminase 1, involucrin, filaggrin, and the cytokeratin pair 1/10 proteins was detected in the conjunctiva of patients with chronic cicatricial phase including Stevens-Johnson syndrome, oculat cicatricial pemphigoid and chemical injuries (134). Keratin 13 (KRT13) was found within the most abundant proteins in the ocular conjunctiva proteome. This data confirmed previous transcriptomic analysis of the human cornea and conjunctiva (135). KRT13 is an acidic keratin expressed in unkeratinized, stratified squamous epithelium (136). This protein has been identified as a specific conjunctival, non-corneal epithelial cell marker using immunocytochemistry applied to impression cytology specimens (137). Moreover, KRT4/KRT13 pair has been used to identify differentiating cells in internal stratified epithelial cells (138). The higher expression of KRT13 in Zikv and Zikv^CZS^ compared to control can be associated to keratinization of conjunctival epithelial cells. Taken together, various keratinization-related proteins were regulated in the comparison between Zikv^CZS^ and Zikv with Ctrl. The molecular data were supported by microscopy analysis. The molecular alterations observed on the ocular surface of infants with CZS were validated by clinical and microscopy analysis. Indeed, the three infants with CZS presented optic disc pallor and excavation and altered macular pigment with pigment deposition. Moreover, bright field microscopy data indicated a higher degree of moderate to severe keratinized ocular epithelial cells in the Zikv^CZS^ compared to control condition (**Figure 6B**).

In order to correlate alteration in the ocular surface proteome with neurodevelopmental abnormalities associated to ZIKV infection, we focused on regulated proteins involved in neurological dysfunctions. Copine-1 (CPNE1) is a phospholipid binding protein located in the plasma membrane of various cell types. In brain cells, CPNE1 is closely associated with AKT signaling pathway, which is important for neural stem cell (NSC) functions during brain development. In this study, abundant expression of CPNE1 was observed in neural lineage cells including NSCs and immature neurons in human (139). Moreover, Copine-1 plays a key role in the activation of mTOR-Akt pathway in neural stem cells during brain development. Over-expression of each C2 domain of CPNE1 increased neurite outgrowth and expression of the neuronal marker protein neurofilament (NF) (139). A fine regulation of Copine-1 expression is crucial for correct brain development. In the ocular surface of infants with CZS the upregulation of Copine-1 indicates the abnormal brain development and could be used as a marker of immature neurons. Similarly, Reticulon-4 subfamily is composed of three proteoforms (Nogo-A, Nogo-B and Nogo-C). RTN4 is an inhibitor of axon outgrowth and causes collapse of neurite growth cone (140–143). Moreover, RTN4A was found to interact with Bcl-2 and Bcl-X_L_, apoptosis-inhibitors, demonstrating a pro-apoptotic role (144). This protein was upregulated in Zikv^CZS^ condition compared to control. Within the same group, Afamin is a vitamin E-binding glycoprotein that is expressed in the liver and is found in plasma, follicular and cerebrospinal fluids (145, 146). Moreover, Afamin was neuroprotective *in vitro* on primary neurons treated with hydrogen peroxide and amyloid (147). Afamin expression was detected in porcine, human postmortem, and mouse brain capillaries and it was identified as α-Tocopherol transport protein across the blood-brain barrier (148). Interestingly, proteomic analysis of the cerebrospinal fluid of rhesus monkeys infected with simian immunodeficiency virus with central nervous system complications revealed downregulation of afamin and vitamin E (148). Due to that, the downregulation of afamin protein on the ocular surface of infants with CZS could be associated to neurodevelopment disorders. PEDF is a secreted 50kDa proteins and a non-inhibitory member of the serine protease inhibitor (SERPIN) gene family with antiangiogenic, anticancer and neurotrophic properties (149, 150). This factor is expressed in all region of the brain and was shown to induce neuronal differentiation in human retinoblastoma cells (149). Moreover, PDEF has been shown to promote neuron survival on cerebellar granule cells, microglia, brain ischemia injury and promote self-renewal of neural stem cells (151–154). The downregulation of PDEF in Zikv^CZS^ compared to Ctrl and Zikv conditions suggests a possible role in neurological diseases.

Impression cytology has been used to detect superficial viral infections such herpes simplex virus, varicella zoster virus, and adenovirus. We searched for Zika virus proteins in the ocular surface proteome and no viral proteins were detected in these samples. However, the presence of ZIKV antigens were detected in the ocular tissue samples from 4 deceased fetuses with a diagnosis of CZS using a ZIKV NS2B protein antibody (21). The antigen was detected in the iris, neural retina and in the optic nerve. Moreover, RT-PCR analysis of the aqueous sample from a Brazilian man with ZIKV infection was positive and ZIKV was identified in conjunctival swabs obtained from 6 patients with ZIKV infection (155, 156). In our data, ZIKV proteins were not detected using the CImP method. This could be due to the eye region sampled and the low abundance of viral proteins compared to host proteins. Moreover, the infants were infected in the first trimester of gestation and at the time of sample collection the viremia could have been drastically decreased.

The comparison between Zikv and Ctrl conditions revealed 166 differentially modulated proteins associated to pathways previously described for the Zikv^CZS^ group (Supplementary Table 3). Interestingly, the fold changes observed for the Zikv vs Ctrl group were lower indicating a mild but significant remodelling of the ocular proteome in infants exposed to ZIKV during the first trimester of gestation. In particular, IFIT3 and IFIH1 were upregulated in the Zikv conditions compared to control. IFIH1 gene encodes a cytoplasmic receptor of the pattern-recognition receptors family that recognizes viral RNA, playing a role in the innate immune response against viral infection (157, 158). The recognition of viral RNA is an ATP-dependent process that induces IFIH1 polymerization and activation of type 1 interferon signalling cascade (159). IFIH1 loss-of-function variants are unable to produce IFN-β and have been reported to restrict the replication of human respiratory syncytial virus and rhinoviruses (160). The range of non-microcephalic anomalies associated to ZIKV infection pose an important clinical and social challenge. The identification of early molecular alterations can improve our understanding of these effects and help in predicting clinical outcome (161).Taken together, these results call for an active surveillance of ophthalmological complications in children exposed to ZIKV during the first trimester of gestation but without CZS.

## Conclusions

The CImP method present here is a simple, robust, sensitive and effective method for the molecular profiling of ocular surface cells. The first human conjunctival ocular proteome is presented here with more than 2000 proteins identified. The method was applied to understand the molecular alterations in children exposed to ZIKV infection during gestation with and without CZS (**Figure 7**). Proteins involved in neutrophil degranulation, neurodevelopment processes and ocular dysfunctions were validated using parallel reaction monitoring confirming the differential expression between the groups. The data presented here show a regulation of neutrophil degranulation and immune cells infiltration, neurodevelopment and ocular dysfunctions. Our study using the CImP method indicates the occurrence of neutrophil and eosinophil infiltration and degranulation in the conjunctival epithelium in children exposed to ZIKV infection. The combination of molecular markers and microscopic evaluation of neutrophils could improve the diagnostic and prognostic of ocular dysfunctions in ZIKV infected children. However, additional evidences are needed to clarify the pathophysiological role and clinical relevance of neutrophils in this disease. The “ocular fingerprint” would be a powerful tool, guiding the physicians towards a clinical decision on the better treatment aiming at a personalized medicine approach for CZS. Congenital Zika syndrome is associated with a wide spectrum of abnormalities that range from mild to severe symptoms (162). These differences can be attributed to the viral strains and load, time of the infection and the different genetic background of the hosts. Due to that, these parameters should be considered to get a deeper understanding of this disease. Another important aspect of ocular pathologies in congenital ZIKV infection will be the discordant clinical outcomes in twin pregnancies (163). Indeed, studying ocular abnormalities in twins with congenital ZIKV infection should help elucidate the mechanism of ZIKV eye infection. The CImP method has the potential to provide detailed cytological and molecular data be applied to several ocular and neurological diseases that involve ocular dysfunction (164–166). The availability of advance of large scale and high-throughput technology will enable the CImP method to be applicable to screen large cohorts. Few biomarker studies applied to human tears have gone from the discovery to the validation phase and none of them has gone into the clinic (167). It is urgent to develop standard operating procedures for assessing specific biomarkers using the CImP method and validate them in multicentre clinical studies (168, 169).

**Figure 7:**
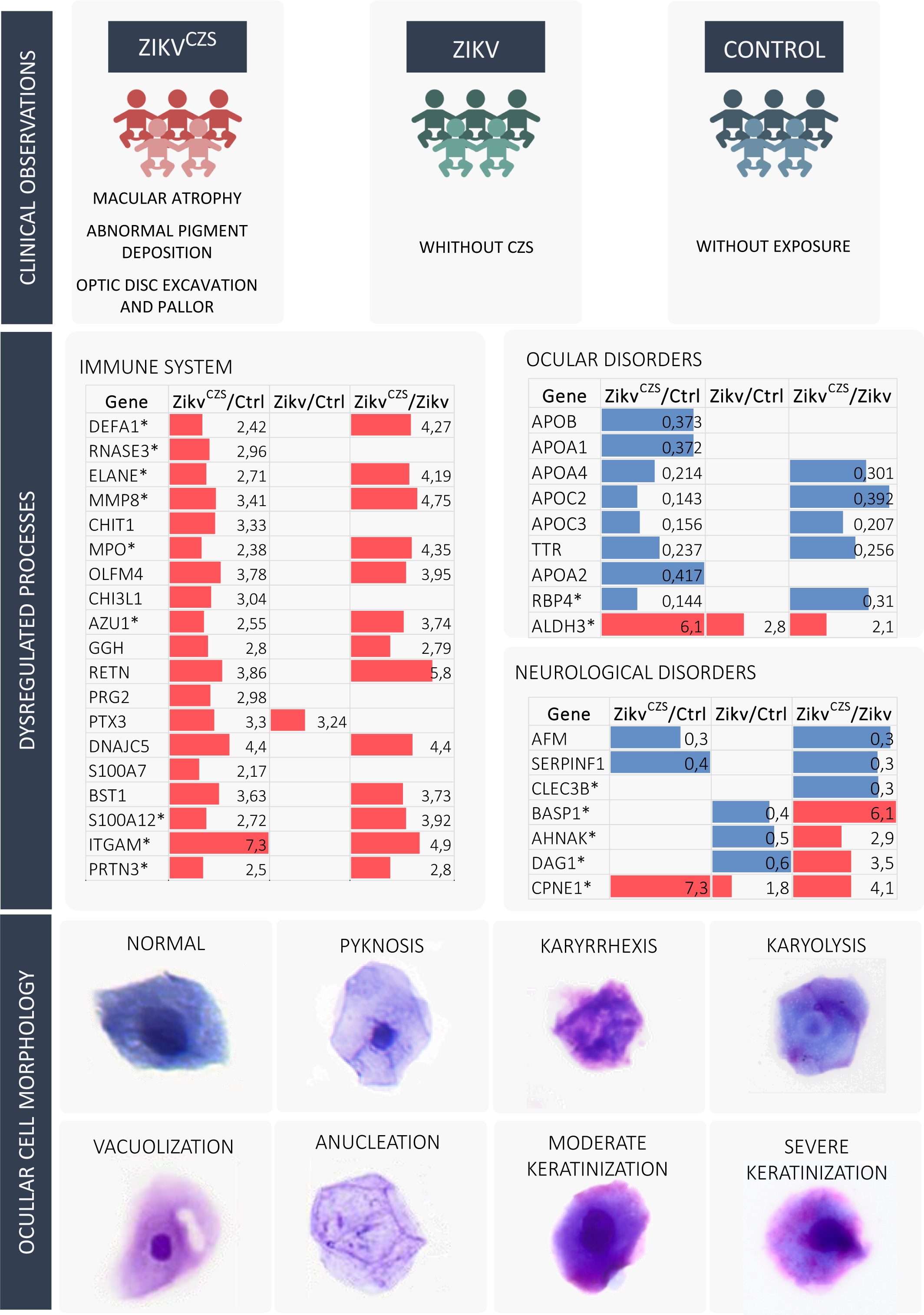
Summary of the findings provided by clinical data, CImP/PRM data, and microscopy showing biological events related to gestational ZIKV exposure with and without CZS. Impression cytology membranes collected from a cohort of infants exposed to ZIKV during the first trimester of gestation and with CZS (Zikv^CZS^) revealed ocular alterations such as optic disc excavation and pallor, abnormal pigment deposition and macular atrophy. These alterations were not detected in the Zikv and Ctrl conditions. The Cellular Imprinting Proteomics approach detected several proteins involved in immune response, cell death, ocular and neurological disorders differentially regulated. These molecular alterations were confirmed by microscopy analysis with defects in the morphology of epithelial cells collected from the Zikv^CZS^ group.

## Author contribution

LR-F, RHB, GP designed the experiments and performed CImP sample preparation. LR-F and GP performed analytical method development, mass spectrometry analysis, data interpretation, PRM analysis and wrote the manuscript. RHB collected the impression cytology membranes and respective informed and written consents. CAC performed the cohort selection and coordinates the clinical follow-up study. MLBS, CBA and GSO assisted on CImP sample preparation. BL assisted in the experimental design. MRL assisted on analytical method development, mass spectrometry analysis and revision of the manuscript. TPS and RCM performed microscopy analyses. All authors contributed in editing the manuscript and approved the final version.

## Data availability

The mass spectrometry-based proteomics data have been deposited to the ProteomeXchange Consortium.

## Acknowledgments

RHB was supported by FAPERJ (201.779/2017), GSO was supported by FAPESP (2018/13283-4), GP was supported by FAPESP (2014/06863-3, 2018/18257-1, 2018/15549-1) and CNPq (bolsa de produtividade), MLBS was supported by CAPES (Finance Code 001), RCM was supported by CNPq (434914/2018-5 and 309734/2018-5). This work was also supported by the VILLUM Center for Bioanalytical Sciences at the University of Southern Denmark.

## Supplementary Figures

**Supplementary Figure S1:**
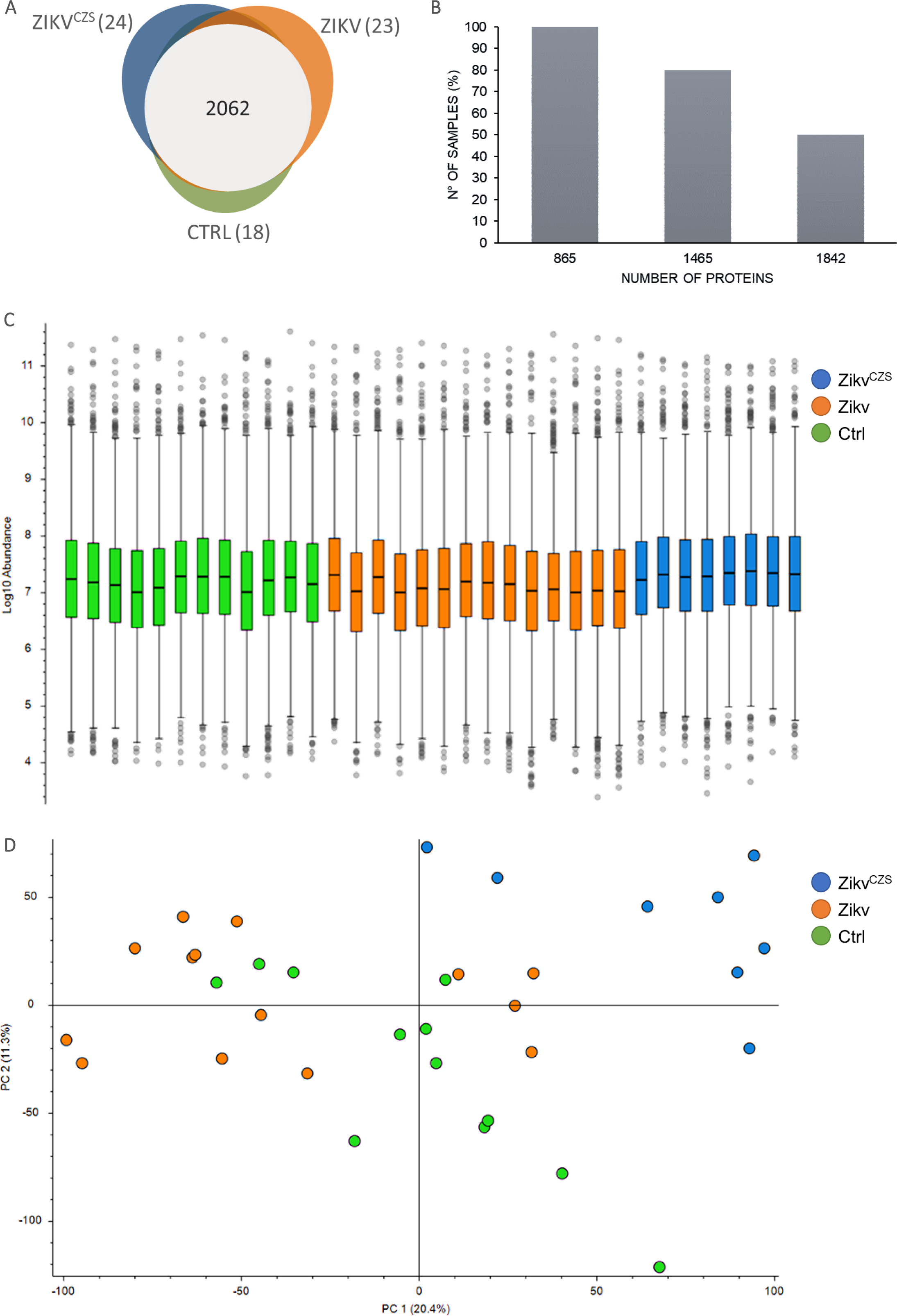
A) Comparison of proteins identified from impression cytology membranes using the CImP method. The membranes were collected from infants exposed to ZIKV during the first trimester with (Zikv^CZS^) and without microcephaly (Zikv) and infants not exposed to ZIKV infection (Ctrl). B) Percentage of proteins identified in 100, 80 and 50% of the samples investigated. C) Distribution of protein abundance of samples within the three conditions. D) Principal component analysis of the samples belonging to the three conditions using the total quantified proteins.

**Supplementary Figure S2:**
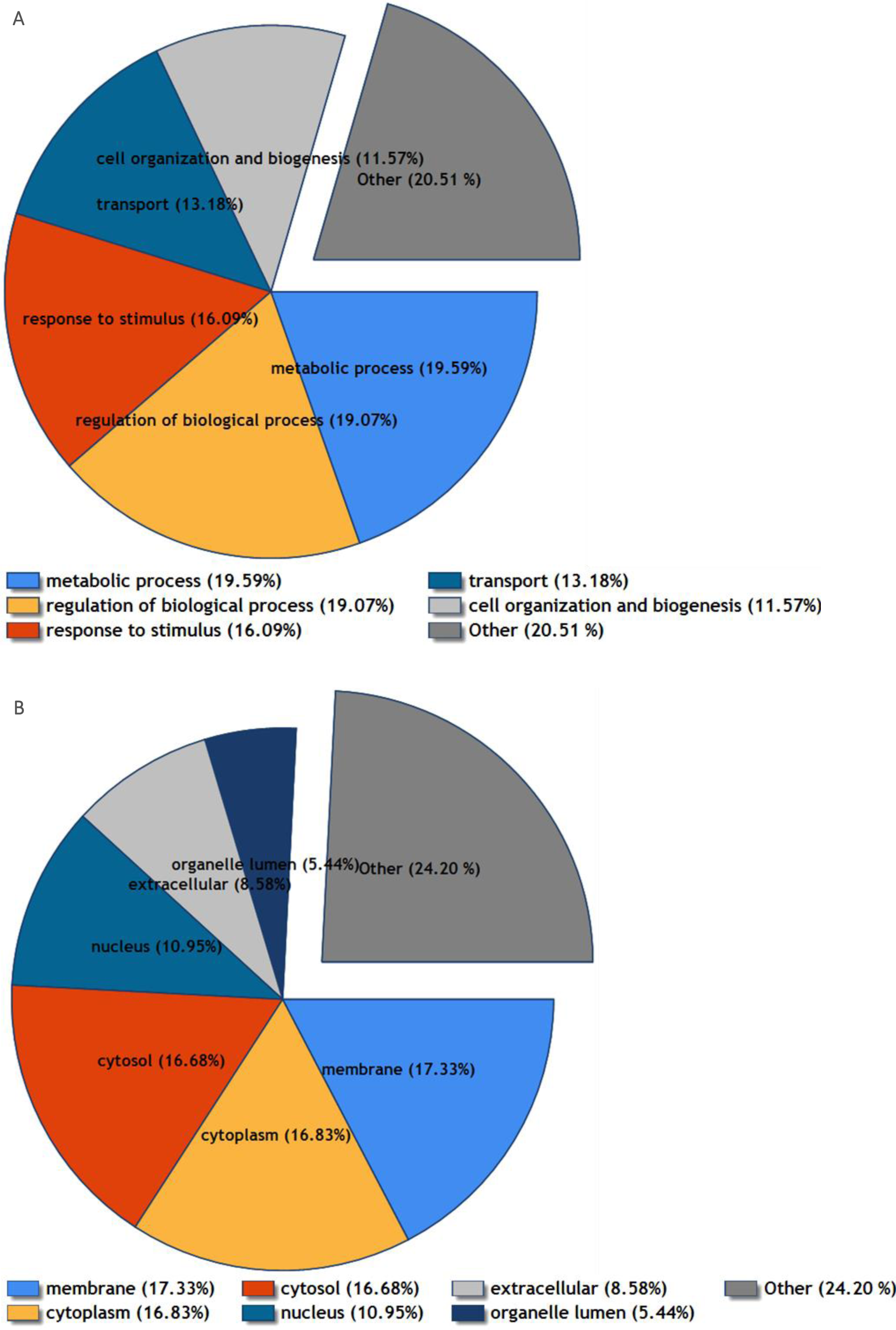
Gene ontology analysis of the total identified proteins using the Proteome Discoverer 2.3 node. A) Biological processes enriched in the total identified proteins. B) Cellular components enriched in the total identified proteins.

**Supplementary Figure S3:**
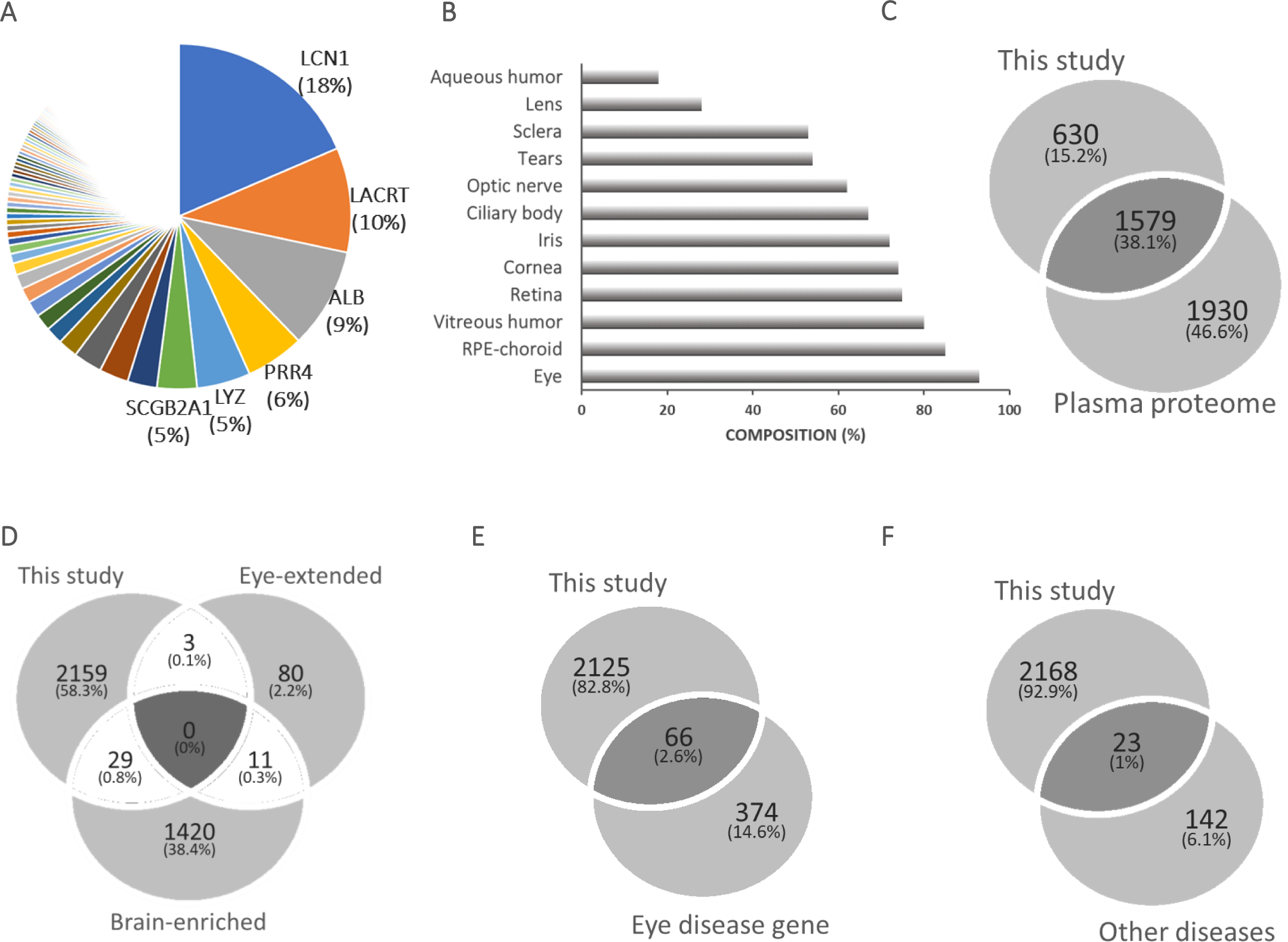
A) Abundance of proteins identified in the epithelial cells and fluid extracted using the impression cytology technique. Intensity-based absolute quantification (iBAQ) was used to calculate the intra-sample protein abundance and the proteins were ranked based on their abundance. B) Comparison of proteins identified using the impression cytology method with the proteome of different ocular tissues and fluids (tears, aqueous humour, lens, vitreous humour, retina, and retinal pigment epithelium (RPE)/choroid, according to Ahmad *et.al.* (59). C) Comparison of proteins identified using the impression cytology method in this study and the human plasma proteome, according to Schwenk *et.al.* (60). D) Comparison of proteins identified using the impression cytology method in this study and brain and eye-enriched proteins reported in the Human Protein Atlas database. E) Comparison of proteins identified using the impression cytology method in this study and genes involved in eye diseases reported in the NEIBank database. F) Comparison of proteins identified using the impression cytology method in this study and genes identified in the eye compartments involved in other diseases as reported in the NEIBank database.

**Supplementary Figure S4:**
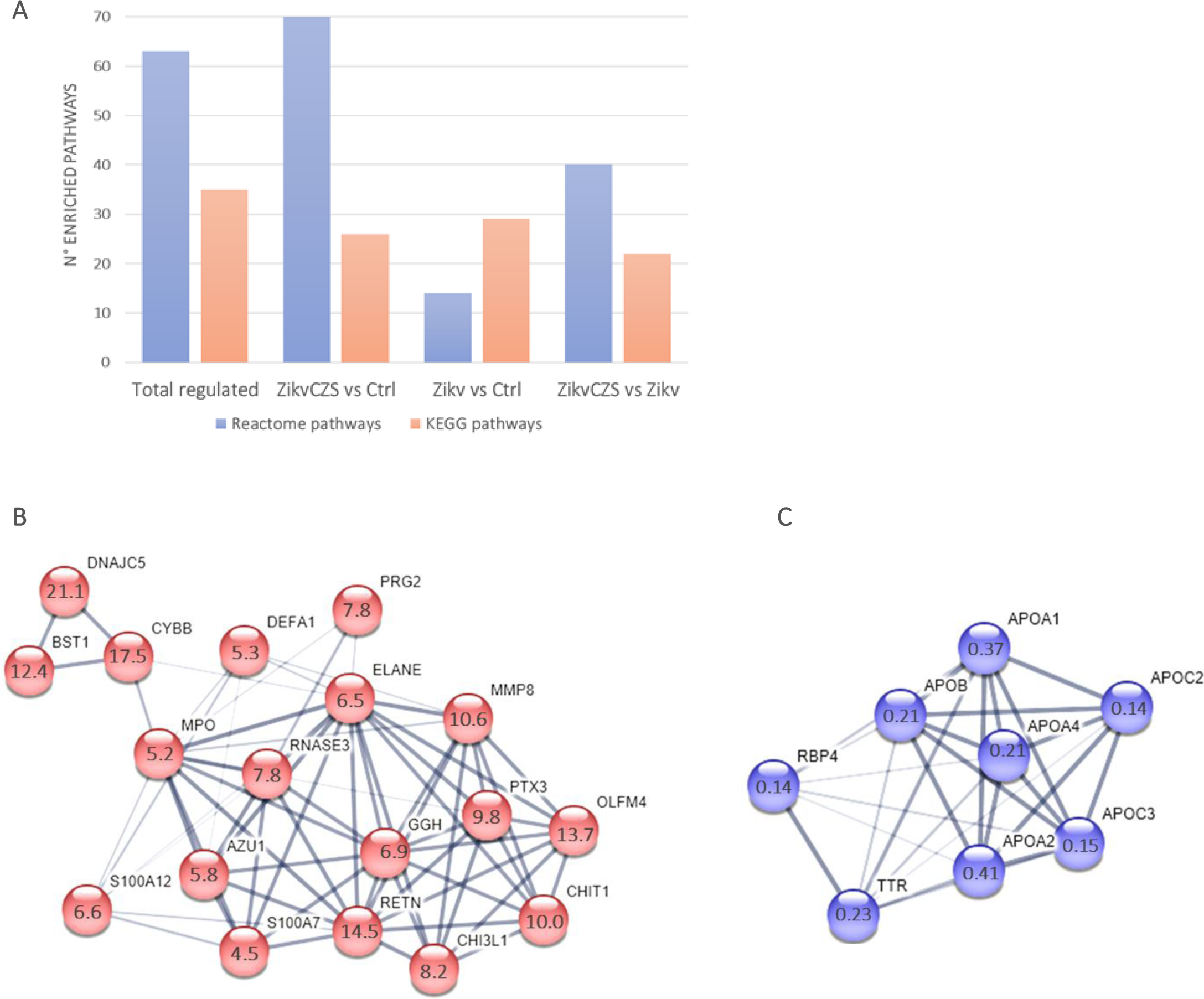
Molecular pathways enriched using the differentially regulated proteins. A) Number of Reactome and KEGG pathways enriched in the differentially regulated proteins datasets (FDR<0.05). B) Protein-protein interaction map of the differentially regulated proteins involved in the immune response pathway in the Zikv^CZS^ vs Ctrl comparison. C) Protein-protein interaction map of the differentially regulated proteins involved in ocular disorders in the Zikv^CZS^ vs Ctrl.

**Supplementary Figure S5:**
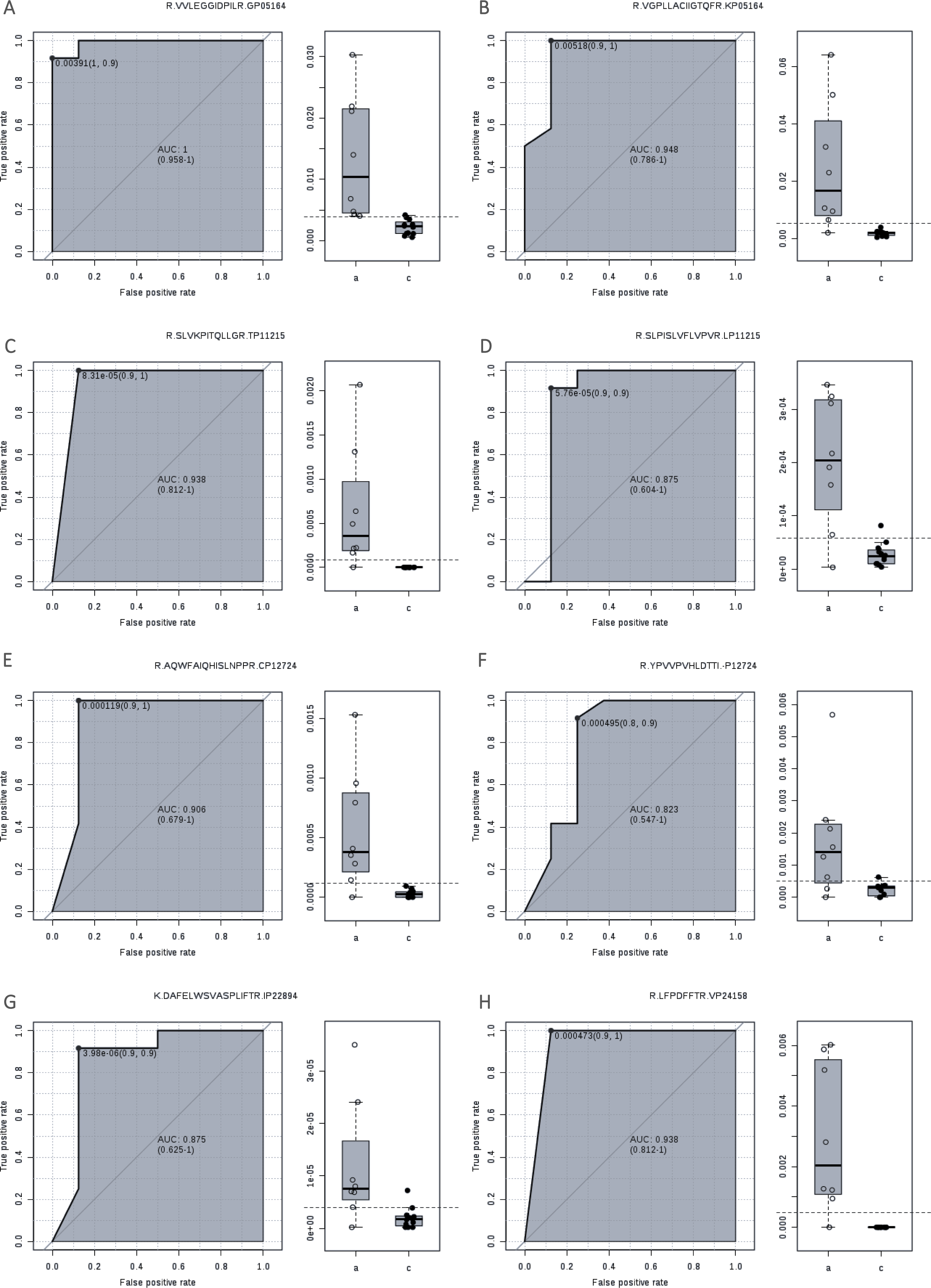
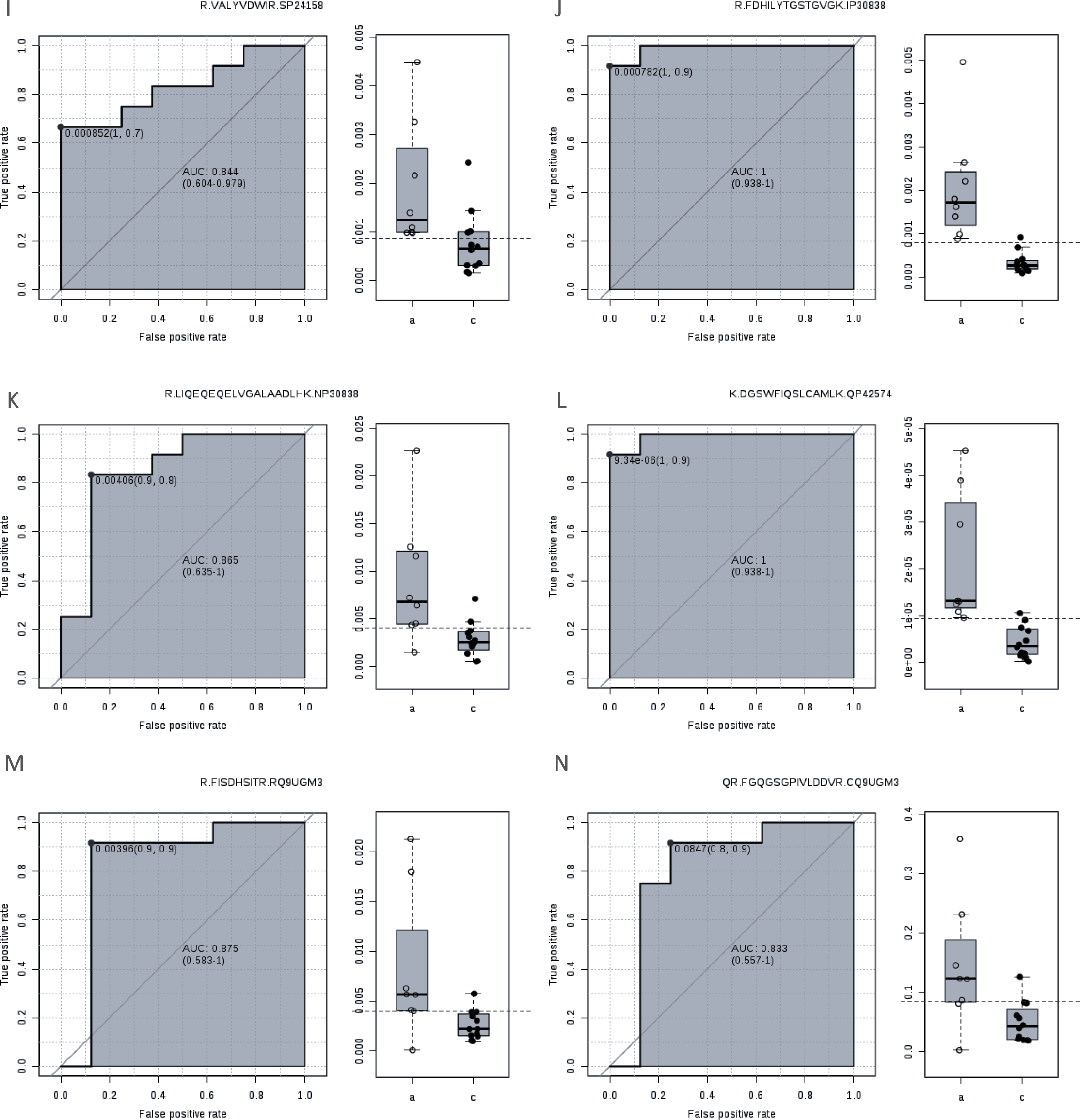
Validation of differentially expressed proteins by PRM in the Zikv^CZS^ vs Ctrl comparison. A-P) Peptides belonging to protein candidates that discriminated the Zikv^CZS^ and Ctrl conditions with AUC>0.8. A,B) P05164 (MPO, Myeloperoxidase); C,D) P11215 (ITGAM, Integrin alpha-M); E,F) P12724 (RNASE3, Eosinophil cationic protein); G) P22894 (MMP8, Neutrophil collagenase); H,I) P24158 (PRTN3, Myeloblastin); J,K) P30838 (ALDH3A1, Aldehyde dehydrogenase, dimeric NADP-preferring); L) P42574 (CASP3, Caspase-3) and M,N) Q9UGM3 (DMBT1, Deleted in malignant brain tumors 1 protein).

**Supplementary Figure S6:**
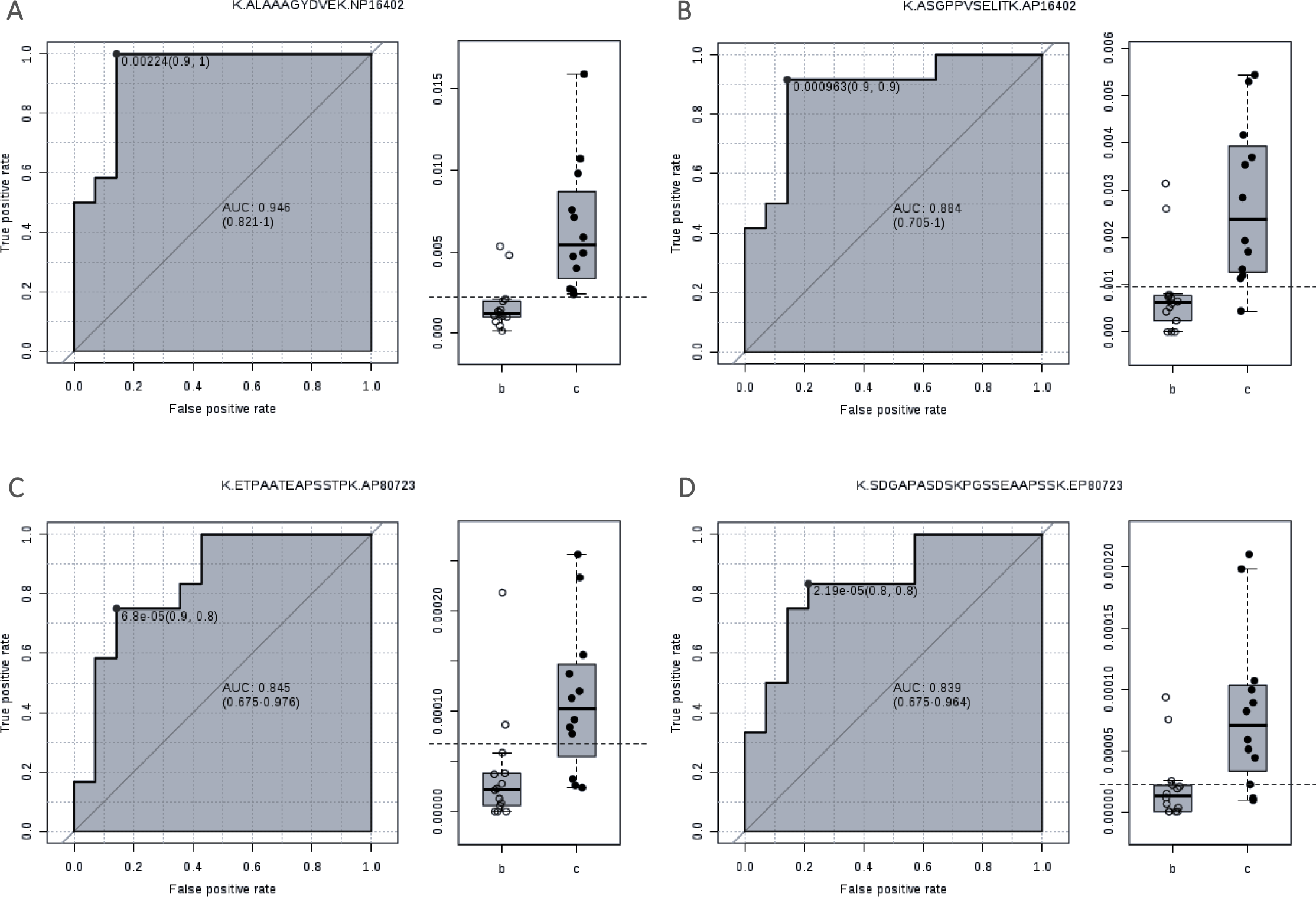
Validation of differentially expressed proteins by PRM in the Zikv vs Ctrl comparison. A-D) Peptides belonging to protein candidates that discriminated the Zikv and Ctrl conditions with AUC>0.8. A,B) P16402 (HIST1H1D, Histone H1.3) and C,D) P80723 (BASP1, Brain acid soluble protein 1).

**Supplementary Figure S7:**
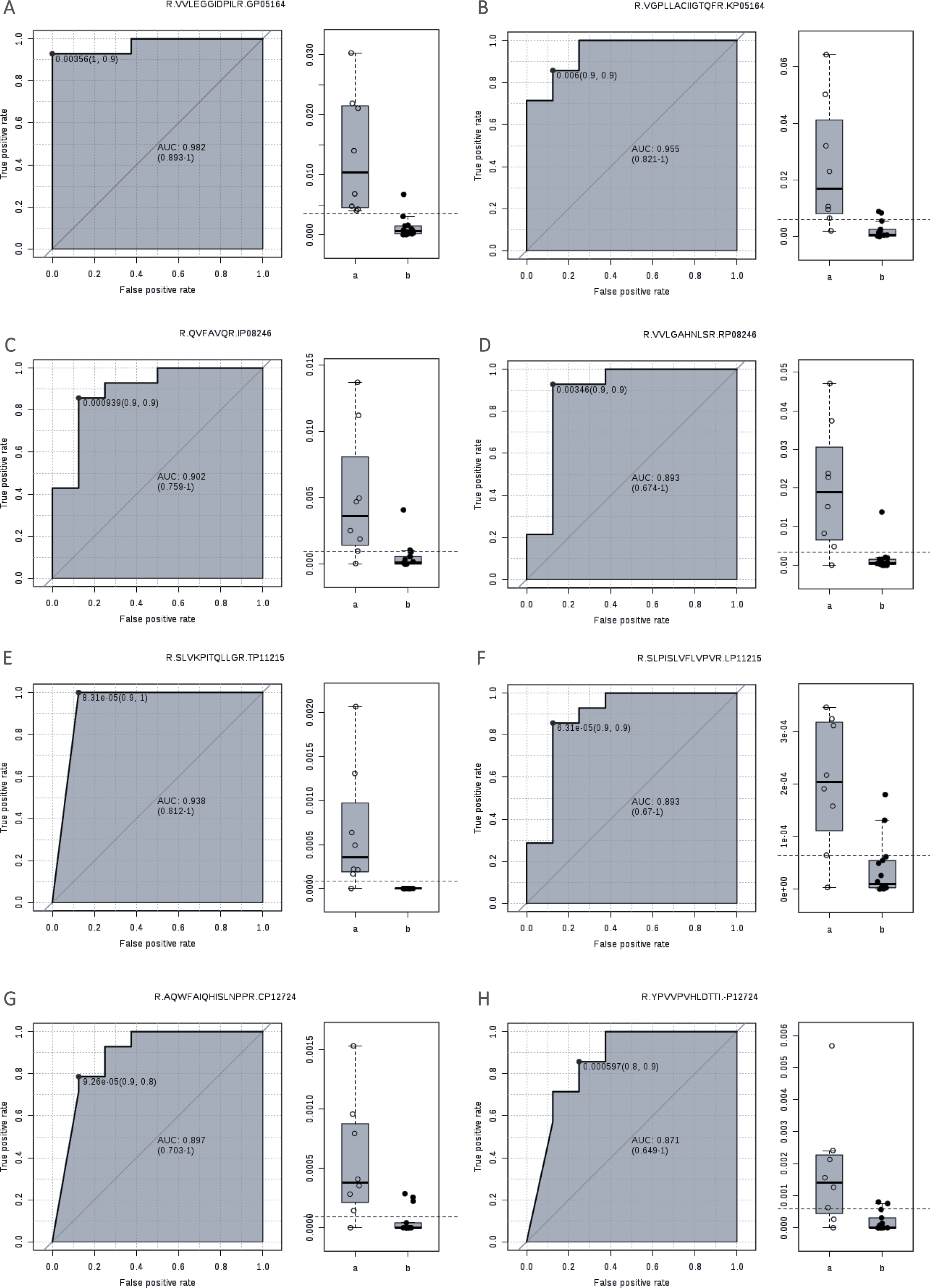
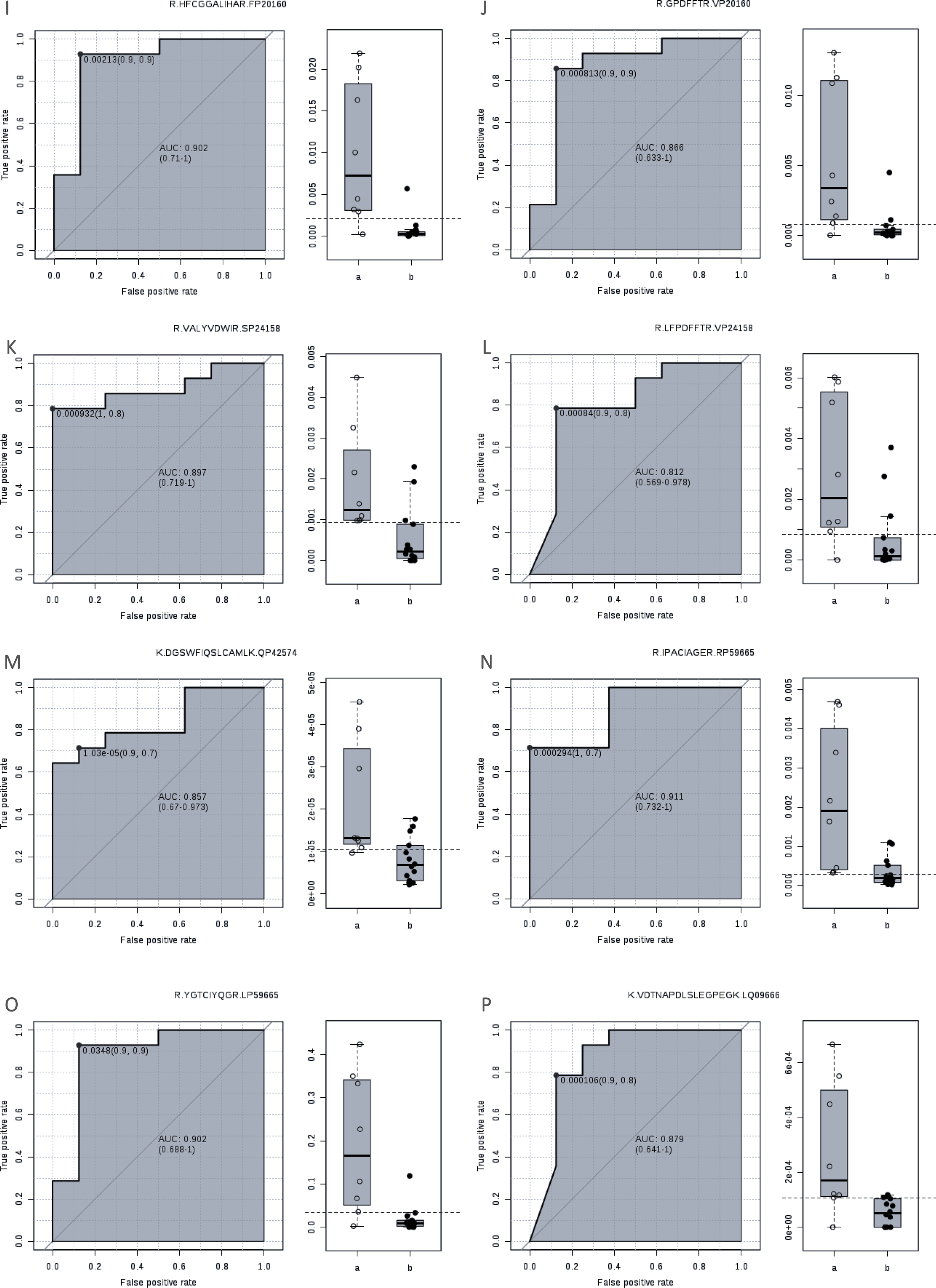
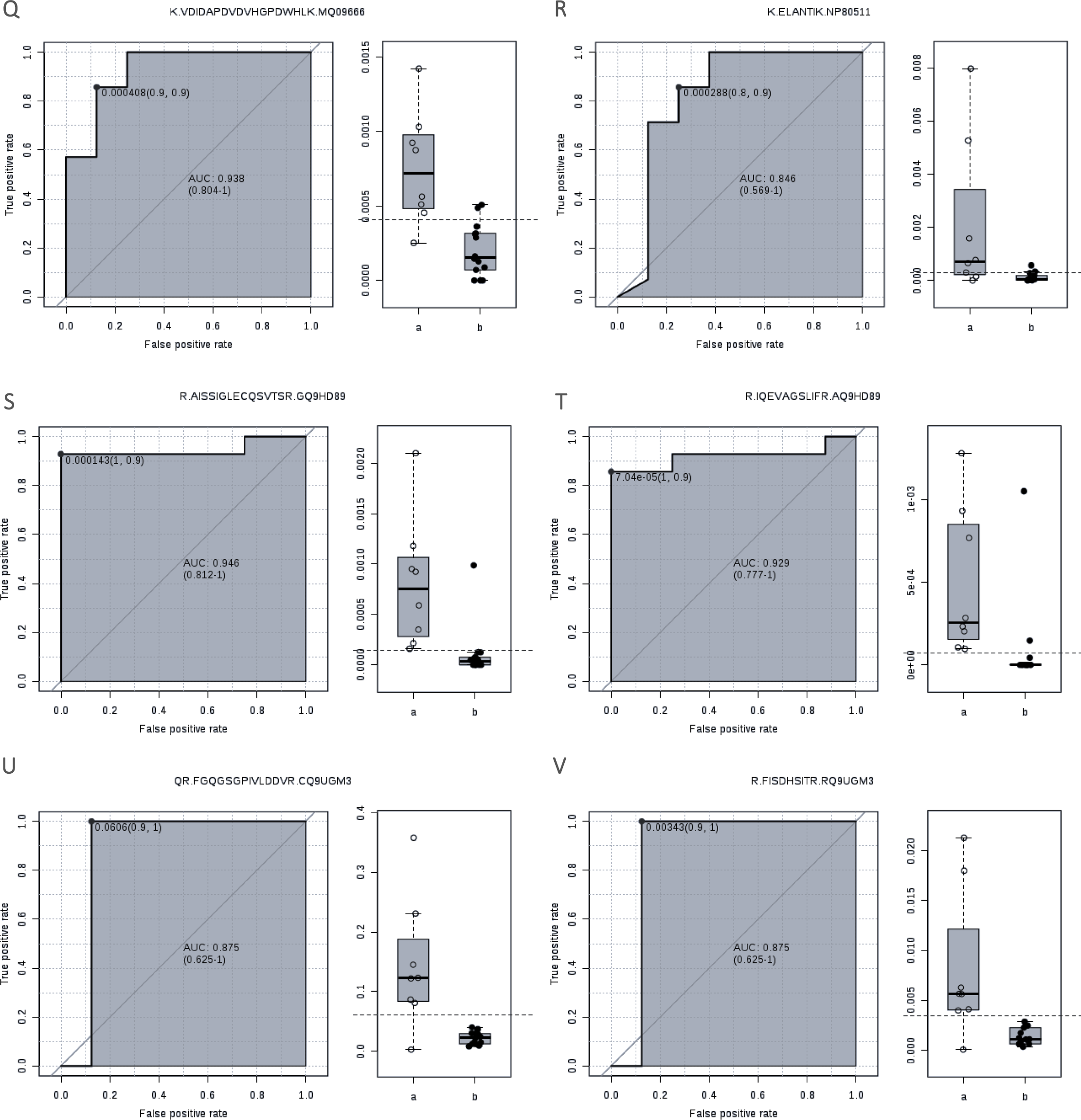
Validation of differentially expressed proteins by PRM in the Zikv^CZS^ vs Zikv comparison. A-Y) Peptides belonging to protein candidates that discriminated the Zikv and Zikv conditions with AUC>0.8. A,B) P05164 (MPO, Myeloperoxidase); C,D) P08246 (ELANE, Neutrophil elastase); E,F) P11215 (ITGAM, Integrin alpha-M); G,H) P12724 (RNASE3, Eosinophil cationic protein); I,J) P20160 (AZU1, Azurocidin); K,L) P24158 (PRTN3, Myeloblastin); M) P42574 (CASP3, Caspase-3); N,O) P59665 (DEFA1, Neutrophil defensin 1); P,Q) Q09666 (AHNAK, Neuroblast differentiation-associated protein AHNAK), R) P80511 (S100A12, Protein S100-A12); S,T) Q9HD89 (RETN, Resistin) and U,V) Q9UGM3 (DMBT1, Deleted in malignant brain tumors 1 protein).

## Supplementary Tables

**Supplementary Table S1:** List of peptide ions targeted for validation. Mass (m/z), charge state (z), peptide sequence and gene names. Cysteine residues in small letter indicate carbamidomethylation modification.

**Supplementary Table S2**: proteins identified and quantified in the total samples. The computational platform Proteome Discoverer 2.3 (Thermo Fisher) was used for protein identification and quantification as reported in the Materials and Methods section. The different columns indicate: Uniprot accession number, Protein description, protein coverage [%], number of Peptides, number of PSMs, number of unique Peptides, Gene Ontology categories, KEGG Pathways, Reactome Pathways, A/C, B/C, A/B, Abundance Ratio Adjusted p-Value: A/C, Abundance Ratio Adjusted p-Value: B/C, Abundance Ratio Adjusted p-Value: A/B, Abundances (normalized) samples A, Abundances (normalized) samples B, Abundances (normalized) samples C and Gene name. A = Zikv^CZS^ condition, B=Zikv condition and C=Ctrl condition.

**Supplementary Table S3**: proteins differentially regulated in the ZikvCZS vs Ctrl comparison. The computational platform Proteome Discoverer 2.3 (Thermo Fisher) was used for protein identification and quantification as reported in the Materials and Methods section. The different columns indicate: Uniprot accession number, Protein description, protein coverage [%], number of Peptides, number of PSMs, number of unique Peptides, Gene Ontology categories, KEGG Pathways, Reactome Pathways, A/C, B/C, A/B, Abundance Ratio Adjusted p-Value: A/C, Abundance Ratio Adjusted p-Value: B/C, Abundance Ratio Adjusted p-Value: A/B, Abundances (normalized) samples A, Abundances (normalized) samples B, Abundances (normalized) samples C and Gene name. A = Zikv^CZS^ condition, B=Zikv condition and C=Ctrl condition

**Supplementary Table S4**: proteins differentially regulated in the Zikv vs Ctrl comparison. The computational platform Proteome Discoverer 2.3 (Thermo Fisher) was used for protein identification and quantification as reported in the Materials and Methods section. The different columns indicate: Uniprot accession number, Protein description, protein coverage [%], number of Peptides, number of PSMs, number of unique Peptides, Gene Ontology categories, KEGG Pathways, Reactome Pathways, A/C, B/C, A/B, Abundance Ratio Adjusted p-Value: A/C, Abundance Ratio Adjusted p-Value: B/C, Abundance Ratio Adjusted p-Value: A/B, Abundances (normalized) samples A, Abundances (normalized) samples B, Abundances (normalized) samples C and Gene name. A = Zikv^CZS^ condition, B=Zikv condition and C=Ctrl condition

**Supplementary Table 5**: proteins differentially regulated in the Zikv^CZS^ vs Zikv comparison. The computational platform Proteome Discoverer 2.3 (Thermo Fisher) was used for protein identification and quantification as reported in the Materials and Methods section. The different columns indicate: Uniprot accession number, Protein description, protein coverage [%], number of Peptides, number of PSMs, number of unique Peptides, Gene Ontology categories, KEGG Pathways, Reactome Pathways, A/C, B/C, A/B, Abundance Ratio Adjusted p-Value: A/C, Abundance Ratio Adjusted p-Value: B/C, Abundance Ratio Adjusted p-Value: A/B, Abundances (normalized) samples A, Abundances (normalized) samples B, Abundances (normalized) samples C and Gene name. A = Zikv^CZS^ condition, B=Zikv condition and C=Ctrl condition

**Supplementary Table 6**: validation of differentially expressed proteins by PRM analysis. Two peptides per protein were monitored by PRM as described in the Materials and Methods section. The raw data were searched in the Proteome discoverer 2.3 (Thermo) computational platform in order to identify and quantify them in all samples.

## References

1. Lee CY, Ng LFP. Zika virus: from an obscurity to a priority. Microbes and infection. 2018;20(11-12):635–45.

2. Musso D, Roche C, Robin E, Nhan T, Teissier A, Cao-Lormeau VM. Potential sexual transmission of Zika virus. Emerging infectious diseases. 2015;21(2):359–61.

3. Dick GW, Kitchen SF, Haddow AJ. Zika virus I. Isolations and serological specificity. Transactions of the Royal Society of Tropical Medicine and Hygiene. 1952;46(5):509–20.

4. Posen HJ, Keystone JS, Gubbay JB, Morris SK. Epidemiology of Zika virus, 1947-2007. BMJ global health. 2016;1(2):e000087.

5. Lanciotti RS, Kosoy OL, Laven JJ, Velez JO, Lambert AJ, Johnson AJ, et al. Genetic and serologic properties of Zika virus associated with an epidemic, Yap State, Micronesia, 2007. Emerging infectious diseases. 2008;14(8):1232–9.

6. de Araujo TVB, Ximenes RAA, Miranda-Filho DB, Souza WV, Montarroyos UR, de Melo APL, et al. Association between microcephaly, Zika virus infection, and other risk factors in Brazil: final report of a case-control study. The Lancet Infectious diseases. 2018;18(3):328–36.

7. Soares de Oliveira-Szejnfeld P, Levine D, Melo AS, Amorim MM, Batista AG, Chimelli L, et al. Congenital Brain Abnormalities and Zika Virus: What the Radiologist Can Expect to See Prenatally and Postnatally. Radiology. 2016;281(1):203–18.

8. Moore CA, Staples JE, Dobyns WB, Pessoa A, Ventura CV, Fonseca EB, et al. Characterizing the Pattern of Anomalies in Congenital Zika Syndrome for Pediatric Clinicians. JAMA pediatrics. 2017;171(3):288–95.

9. Martinot AJ, Abbink P, Afacan O, Prohl AK, Bronson R, Hecht JL, et al. Fetal Neuropathology in Zika Virus-Infected Pregnant Female Rhesus Monkeys. Cell. 2018;173(5):1111–22 e10.

10. Miner JJ, Sene A, Richner JM, Smith AM, Santeford A, Ban N, et al. Zika Virus Infection in Mice Causes Panuveitis with Shedding of Virus in Tears. Cell reports. 2016;16(12):3208–18.

11. Mohr EL, Block LN, Newman CM, Stewart LM, Koenig M, Semler M, et al. Ocular and uteroplacental pathology in a macaque pregnancy with congenital Zika virus infection. PloS one. 2018;13(1):e0190617.

12. Nguyen SM, Antony KM, Dudley DM, Kohn S, Simmons HA, Wolfe B, et al. Highly efficient maternal-fetal Zika virus transmission in pregnant rhesus macaques. PLoS pathogens. 2017;13(5):e1006378.

13. Sahiner F, Sig AK, Savasci U, Tekin K, Akay F. Zika Virus-associated Ocular and Neurologic Disorders: The Emergence of New Evidence. The Pediatric infectious disease journal. 2017;36(12):e341–e6.

14. de Paula Freitas B, de Oliveira Dias JR, Prazeres J, Sacramento GA, Ko AI, Maia M, et al. Ocular Findings in Infants With Microcephaly Associated With Presumed Zika Virus Congenital Infection in Salvador, Brazil. JAMA ophthalmology. 2016;134(5):529–35.

15. Miranda HA, 2nd, Costa MC, Frazao MAM, Simao N, Franchischini S, Moshfeghi DM. Expanded Spectrum of Congenital Ocular Findings in Microcephaly with Presumed Zika Infection. Ophthalmology. 2016;123(8):1788–94.

16. de Oliveira Dias JR, Ventura CV, Borba PD, de Paula Freitas B, Pierroti LC, do Nascimento AP, et al. Infants with Congenital Zika Syndrome and Ocular Findings from Sao Paulo, Brazil: Spread of Infection. Retinal cases & brief reports. 2018;12(4):382–6.

17. Ventura CV, Maia M, Bravo-Filho V, Gois AL, Belfort R, Jr. Zika virus in Brazil and macular atrophy in a child with microcephaly. Lancet. 2016;387(10015):228.

18. Ventura CV, Ventura LO, Bravo-Filho V, Martins TT, Berrocal AM, Gois AL, et al. Optical Coherence Tomography of Retinal Lesions in Infants With Congenital Zika Syndrome. JAMA ophthalmology. 2016;134(12):1420–7.

19. Guevara JG, Agarwal-Sinha S. Ocular abnormalities in congenital Zika syndrome: a case report, and review of the literature. Journal of medical case reports. 2018;12(1):161.

20. de Paula Freitas B, Ventura CV, Maia M, Belfort R, Jr. Zika virus and the eye. Current opinion in ophthalmology. 2017;28(6):595–9.

21. Fernandez MP, Parra Saad E, Ospina Martinez M, Corchuelo S, Mercado Reyes M, Herrera MJ, et al. Ocular Histopathologic Features of Congenital Zika Syndrome. JAMA ophthalmology. 2017;135(11):1163–9.

22. Ritter JM, Martines RB, Zaki SR. Zika Virus: Pathology From the Pandemic. Archives of pathology & laboratory medicine. 2017;141(1):49–59.

23. Ventura CV, Maia M, Travassos SB, Martins TT, Patriota F, Nunes ME, et al. Risk Factors Associated With the Ophthalmoscopic Findings Identified in Infants With Presumed Zika Virus Congenital Infection. JAMA ophthalmology. 2016;134(8):912–8.

24. Zin AA, Tsui I, Rossetto J, Vasconcelos Z, Adachi K, Valderramos S, et al. Screening Criteria for Ophthalmic Manifestations of Congenital Zika Virus Infection. JAMA pediatrics. 2017;171(9):847–54.

25. Valentine G, Marquez L, Pammi M. Zika Virus-Associated Microcephaly and Eye Lesions in the Newborn. Journal of the Pediatric Infectious Diseases Society. 2016;5(3):323–8.

26. Singh R, Joseph A, Umapathy T, Tint NL, Dua HS. Impression cytology of the ocular surface. The British journal of ophthalmology. 2005;89(12):1655–9.

27. Egbert PR, Lauber S, Maurice DM. A simple conjunctival biopsy. American journal of ophthalmology. 1977;84(6):798–801.

28. Nelson JD, Havener VR, Cameron JD. Cellulose acetate impressions of the ocular surface. Dry eye states. Arch Ophthalmol. 1983;101(12):1869–72.

29. Hagan S. Biomarkers of ocular surface disease using impression cytology. Biomarkers in medicine. 2017;11(12):1135–47.

30. Lopin E, Deveney T, Asbell PA. Impression cytology: recent advances and applications in dry eye disease. The ocular surface. 2009;7(2):93–110.

31. Brignole-Baudouin F, Riancho L, Ismail D, Deniaud M, Amrane M, Baudouin C. Correlation Between the Inflammatory Marker HLA-DR and Signs and Symptoms in Moderate to Severe Dry Eye Disease. Investigative ophthalmology & visual science. 2017;58(4):2438–48.

32. Nicolle P, Liang H, Reboussin E, Rabut G, Warcoin E, Brignole-Baudouin F, et al. Proinflammatory Markers, Chemokines, and Enkephalin in Patients Suffering from Dry Eye Disease. International journal of molecular sciences. 2018;19(4).

33. Aronni S, Cortes M, Sacchetti M, Lambiase A, Micera A, Sgrulletta R, et al. Upregulation of ICAM-1 expression in the conjunctiva of patients with chronic graft-versus-host disease. European journal of ophthalmology. 2006;16(1):17–23.

34. Corrales RM, Narayanan S, Fernandez I, Mayo A, Galarreta DJ, Fuentes-Paez G, et al. Ocular mucin gene expression levels as biomarkers for the diagnosis of dry eye syndrome. Investigative ophthalmology & visual science. 2011;52(11):8363–9.

35. Wittpenn JR, Tseng SC, Sommer A. Detection of early xerophthalmia by impression cytology. Arch Ophthalmol. 1986;104(2):237–9.

36. Karalezli A, Borazan M, Altinors DD, Dursun R, Kiyici H, Akova YA. Conjunctival impression cytology, ocular surface, and tear-film changes in patients treated with systemic isotretinoin. Cornea. 2009;28(1):46–50.

37. Vanathi M, Kashyap S, Khan R, Seth T, Mishra P, Mahapatra M, et al. Ocular surface evaluation in allogenic hematopoietic stem cell transplantation patients. European journal of ophthalmology. 2014;24(5):655–66.

38. Okada N, Fukagawa K, Takano Y, Dogru M, Tsubota K, Fujishima H, et al. The implications of the upregulation of ICAM-1/VCAM-1 expression of corneal fibroblasts on the pathogenesis of allergic keratopathy. Investigative ophthalmology & visual science. 2005;46(12):4512–8.

39. Shoji J, Inada N, Sawa M. Evaluation of eotaxin-1, -2, and -3 protein production and messenger RNA expression in patients with vernal keratoconjunctivitis. Japanese journal of ophthalmology. 2009;53(2):92–9.

40. Shiraki Y, Shoji J, Inada N. Clinical Usefulness of Monitoring Expression Levels of CCL24 (Eotaxin-2) mRNA on the Ocular Surface in Patients with Vernal Keratoconjunctivitis and Atopic Keratoconjunctivitis. Journal of ophthalmology. 2016;2016:3573142.

41. Citirik M, Berker N, Haksever H, Elgin U, Ustun H. Conjunctival impression cytology in non-proliferative and proliferative diabetic retinopathy. International journal of ophthalmology. 2014;7(2):321–5.

42. Yoon KC, Im SK, Seo MS. Changes of tear film and ocular surface in diabetes mellitus. Korean journal of ophthalmology : KJO. 2004;18(2):168–74.

43. Hsu SL, Lee PY, Chang CH, Chen CH. Immunological impression cytology of the conjunctival epithelium in patients with thyroid orbitopathy-related dry eye. Genetics and molecular research : GMR. 2016;15(3).

44. Uzel MM, Citirik M, Kekilli M, Cicek P. Local ocular surface parameters in patients with systemic celiac disease. Eye (Lond). 2017;31(7):1093–8.

45. Mrugacz M, Zak J, Bakunowicz-Lazarczyk A, Wysocka J, Minarowska A. Flow cytometric analysis of HLA-DR antigen in conjunctival epithelial cells of patients with cystic fibrosis. Eye (Lond). 2007;21(8):1062–6.

46. Andrade FEC, Correa MP, Gimenes AD, Dos Santos MS, Campos M, Chammas R, et al. Galectin-3: role in ocular allergy and potential as a predictive biomarker. The British journal of ophthalmology. 2018;102(7):1003–10.

47. Dachir S, Gutman H, Gore A, Cohen L, Cohen M, Amir A, et al. Ocular Surface Changes After Sulfur Mustard Exposure in Rabbits, Monitored by Impression Cytology. Cornea. 2017;36(8):980–7.

48. Chahota R, Ogawa H, Ohya K, Yamaguchi T, Everett KDE, Fukushi H. Involvement of multiple Chlamydia suis genotypes in porcine conjunctivitis. Transboundary and emerging diseases. 2018;65(1):272–7.

49. Doughty MJ. Objective assessment of conjunctival squamous metaplasia by measures of cell and nucleus dimensions. Diagnostic cytopathology. 2011;39(6):409–23.

50. Danjo Y, Watanabe H, Tisdale AS, George M, Tsumura T, Abelson MB, et al. Alteration of mucin in human conjunctival epithelia in dry eye. Investigative ophthalmology & visual science. 1998;39(13):2602–9.

51. Lopez-Miguel A, Gutierrez-Gutierrez S, Garcia-Vazquez C, Enriquez-de-Salamanca A. RNA Collection From Human Conjunctival Epithelial Cells Obtained With a New Device for Impression Cytology. Cornea. 2017;36(1):59–63.

52. Soria J, Acera A, Duran JA, Boto-de-Los-Bueis A, Del-Hierro-Zarzuelo A, Gonzalez N, et al. The analysis of human conjunctival epithelium proteome in ocular surface diseases using impression cytology and 2D-DIGE. Experimental eye research. 2018;167:31-43.

53. Raquel Hora Barbosa MLBdS, Thiago P. Silva, Liva Rosa-Fernandes, Ana Pinto, Pricila S. Spínola, Cibele R. Bonvicino, Priscila V. Fernandes, Evandro Lucena, Giuseppe Palmisano, Rossana C. N. Melo, Claudete Cardoso, Bernardo Lemos. Impression cytology is a non-invasive and effective method for ocular cell obtention from babies with Congenital Zika Syndrome: perspectives in OMIC studies. BioRxiv. 2019.

54. Vianna RAO, Lovero KL, Oliveira SA, Fernandes AR, Santos T, Lima L, et al. Children Born to Mothers with Rash During Zika Virus Epidemic in Brazil: First 18 Months of Life. J Trop Pediatr. 2019.

55. Brasil, Ministerio da Saude. Orientações integradas de vigilância e atenção à saúde no âmbito da Emergência de Saúde Pública de Importância Nacional 2017. Available from: http://bvsms.saude.gov.br/publicacoes/orientacoes_emergencia_gestacao_infancia_zika.pdf

56. Kawahara R, Rosa-Fernandes L, Dos Santos AF, Bandeira CL, Dombrowski JG, Souza RM, et al. Integrated Proteomics Reveals Apoptosis-related Mechanisms Associated with Placental Malaria. Mol Cell Proteomics. 2019;18(2):182–99.

57. Gobom J, Nordhoff E, Mirgorodskaya E, Ekman R, Roepstorff P. Sample purification and preparation technique based on nano-scale reversed-phase columns for the sensitive analysis of complex peptide mixtures by matrix-assisted laser desorption/ionization mass spectrometry. J Mass Spectrom. 1999;34(2):105–16.

58. Schwanhausser B, Busse D, Li N, Dittmar G, Schuchhardt J, Wolf J, et al. Global quantification of mammalian gene expression control. Nature. 2011;473(7347):337–42.

59. Ahmad MT, Zhang P, Dufresne C, Ferrucci L, Semba RD. The Human Eye Proteome Project: Updates on an Emerging Proteome. Proteomics. 2018;18(5-6):e1700394.

60. Schwenk JM, Omenn GS, Sun Z, Campbell DS, Baker MS, Overall CM, et al. The Human Plasma Proteome Draft of 2017: Building on the Human Plasma PeptideAtlas from Mass Spectrometry and Complementary Assays. J Proteome Res. 2017;16(12):4299–310.

61. Singh MS, Marquezan MC, Omiadze R, Reddy AK, Belfort R, Jr., May WN. Inner retinal vasculopathy in Zika virus disease. Am J Ophthalmol Case Rep. 2018;10:6-7.

62. Glasgow BJ, Abduragimov AR, Farahbakhsh ZT, Faull KF, Hubbell WL. Tear lipocalins bind a broad array of lipid ligands. Curr Eye Res. 1995;14(5):363–72.

63. Elangovan N, Lee YC, Tzeng WF, Chu ST. Delivery of ferric ion to mouse spermatozoa is mediated by lipocalin internalization. Biochem Biophys Res Commun. 2004;319(4):1096–104.

64. Kjeldsen L, Cowland JB, Borregaard N. Human neutrophil gelatinase-associated lipocalin and homologous proteins in rat and mouse. Biochim Biophys Acta. 2000;1482(1-2):272–83.

65. Flo TH, Smith KD, Sato S, Rodriguez DJ, Holmes MA, Strong RK, et al. Lipocalin 2 mediates an innate immune response to bacterial infection by sequestrating iron. Nature. 2004;432(7019):917–21.

66. Moschen AR, Adolph TE, Gerner RR, Wieser V, Tilg H. Lipocalin-2: A Master Mediator of Intestinal and Metabolic Inflammation. Trends Endocrinol Metab. 2017;28(5):388–97.

67. Dota A, Nishida K, Adachi W, Nakamura T, Koizumi N, Kawamoto S, et al. An expression profile of active genes in human conjunctival epithelium. Experimental eye research. 2001;72(3):235–41.

68. Stoeckelhuber M, Messmer EM, Schmidt C, Xiao F, Schubert C, Klug J. Immunohistochemical analysis of secretoglobin SCGB 2A1 expression in human ocular glands and tissues. Histochem Cell Biol. 2006;126(1):103–9.

69. McDermott AM. Antimicrobial compounds in tears. Experimental eye research. 2013;117:53-61.

70. Bai F, Kong KF, Dai J, Qian F, Zhang L, Brown CR, et al. A paradoxical role for neutrophils in the pathogenesis of West Nile virus. J Infect Dis. 2010;202(12):1804–12.

71. Srivastava S, Khanna N, Saxena SK, Singh A, Mathur A, Dhole TN. Degradation of Japanese encephalitis virus by neutrophils. Int J Exp Pathol. 1999;80(1):17–24.

72. Jarczak J, Kosciuczuk EM, Lisowski P, Strzalkowska N, Jozwik A, Horbanczuk J, et al. Defensins: natural component of human innate immunity. Hum Immunol. 2013;74(9):1069–79.

73. Daher KA, Selsted ME, Lehrer RI. Direct inactivation of viruses by human granulocyte defensins. J Virol. 1986;60(3):1068–74.

74. Fraisier C, Papa A, Granjeaud S, Hintzen R, Martina B, Camoin L, et al. Cerebrospinal fluid biomarker candidates associated with human WNV neuroinvasive disease. PloS one. 2014;9(4):e93637.

75. Loke P, Hammond SN, Leung JM, Kim CC, Batra S, Rocha C, et al. Gene expression patterns of dengue virus-infected children from nicaragua reveal a distinct signature of increased metabolism. PLoS Negl Trop Dis. 2010;4(6):e710.

76. Amaral DC, Rachid MA, Vilela MC, Campos RD, Ferreira GP, Rodrigues DH, et al. Intracerebral infection with dengue-3 virus induces meningoencephalitis and behavioral changes that precede lethality in mice. J Neuroinflammation. 2011;8:23.

77. Ubol S, Masrinoul P, Chaijaruwanich J, Kalayanarooj S, Charoensirisuthikul T, Kasisith J. Differences in global gene expression in peripheral blood mononuclear cells indicate a significant role of the innate responses in progression of dengue fever but not dengue hemorrhagic fever. J Infect Dis. 2008;197(10):1459–67.

78. Chen HR, Chao CH, Liu CC, Ho TS, Tsai HP, Perng GC, et al. Macrophage migration inhibitory factor is critical for dengue NS1-induced endothelial glycocalyx degradation and hyperpermeability. PLoS pathogens. 2018;14(4):e1007033.

79. Rosenbauer F, Wagner K, Zhang P, Knobeloch KP, Iwama A, Tenen DG. pDP4, a novel glycoprotein secreted by mature granulocytes, is regulated by transcription factor PU.1. Blood. 2004;103(11):4294–301.

80. Zhang J, Liu WL, Tang DC, Chen L, Wang M, Pack SD, et al. Identification and characterization of a novel member of olfactomedin-related protein family, hGC-1, expressed during myeloid lineage development. Gene. 2002;283(1-2):83–93.

81. Brand HK, Ahout IM, de Ridder D, van Diepen A, Li Y, Zaalberg M, et al. Olfactomedin 4 Serves as a Marker for Disease Severity in Pediatric Respiratory Syncytial Virus (RSV) Infection. PloS one. 2015;10(7):e0131927.

82. Liu W, Yan M, Liu Y, McLeish KR, Coleman WG, Jr., Rodgers GP. Olfactomedin 4 inhibits cathepsin C-mediated protease activities, thereby modulating neutrophil killing of Staphylococcus aureus and Escherichia coli in mice. J Immunol. 2012;189(5):2460–7.

83. Liu W, Yan M, Liu Y, Wang R, Li C, Deng C, et al. Olfactomedin 4 down-regulates innate immunity against Helicobacter pylori infection. Proc Natl Acad Sci U S A. 2010;107(24):11056–61.

84. Jampol LM, Ferris FL, 3rd, Bishop RJ. Ebola and the eye. JAMA ophthalmology. 2015;133(10):1105–6.

85. Yahia SB, Khairallah M. Ocular manifestations of West Nile virus infection. Int J Med Sci. 2009;6(3):114–5.

86. Manangeeswaran M, Kielczewski JL, Sen HN, Xu BC, Ireland DDC, McWilliams IL, et al. ZIKA virus infection causes persistent chorioretinal lesions. Emerg Microbes Infect. 2018;7(1):96.

87. Kodati S, Palmore TN, Spellman FA, Cunningham D, Weistrop B, Sen HN. Bilateral posterior uveitis associated with Zika virus infection. Lancet. 2017;389(10064):125–6.

88. Perez VL, Caspi RR. Immune mechanisms in inflammatory and degenerative eye disease. Trends Immunol. 2015;36(6):354–63.

89. Wang W, Li G, De W, Luo Z, Pan P, Tian M, et al. Zika virus infection induces host inflammatory responses by facilitating NLRP3 inflammasome assembly and interleukin-1beta secretion. Nat Commun. 2018;9(1):106.

90. Williams GP, Nightingale P, Southworth S, Denniston AK, Tomlins PJ, Turner S, et al. Conjunctival Neutrophils Predict Progressive Scarring in Ocular Mucous Membrane Pemphigoid. Investigative ophthalmology & visual science. 2016;57(13):5457–69.

91. Williams GP, Tomlins PJ, Denniston AK, Southworth HS, Sreekantham S, Curnow SJ, et al. Elevation of conjunctival epithelial CD45INTCD11b(+)CD16(+)CD14(-) neutrophils in ocular Stevens-Johnson syndrome and toxic epidermal necrolysis. Investigative ophthalmology & visual science. 2013;54(7):4578–85.

92. Heiligenhaus A, Schaller J, Mauss S, Engelbrecht S, Dutt JE, Foster CS, et al. Eosinophil granule proteins expressed in ocular cicatricial pemphigoid. The British journal of ophthalmology. 1998;82(3):312–7.

93. Trocme SD, Bartley GB, Campbell RJ, Gleich GJ, Leiferman KM. Eosinophil and neutrophil degranulation in ophthalmic lesions of Wegener’s granulomatosis. Arch Ophthalmol. 1991;109(11):1585–9.

94. Bowen JR, Quicke KM, Maddur MS, O’Neal JT, McDonald CE, Fedorova NB, et al. Zika Virus Antagonizes Type I Interferon Responses during Infection of Human Dendritic Cells. PLoS pathogens. 2017;13(2):e1006164.

95. Matusali G, Houzet L, Satie AP, Mahe D, Aubry F, Couderc T, et al. Zika virus infects human testicular tissue and germ cells. J Clin Invest. 2018;128(10):4697–710.

96. Wacher C, Muller M, Hofer MJ, Getts DR, Zabaras R, Ousman SS, et al. Coordinated regulation and widespread cellular expression of interferon-stimulated genes (ISG) ISG-49, ISG-54, and ISG-56 in the central nervous system after infection with distinct viruses. J Virol. 2007;81(2):860–71.

97. Kovacs SB, Miao EA. Gasdermins: Effectors of Pyroptosis. Trends Cell Biol. 2017;27(9):673–84.

98. Man SM, Kanneganti TD. Gasdermin D: the long-awaited executioner of pyroptosis. Cell Res. 2015;25(11):1183–4.

99. Gao J, Cui JZ, To E, Cao S, Matsubara JA. Evidence for the activation of pyroptotic and apoptotic pathways in RPE cells associated with NLRP3 inflammasome in the rodent eye. J Neuroinflammation. 2018;15(1):15.

100. Sun N, Zhang H. Pyroptosis in pterygium pathogenesis. Biosci Rep. 2018;38(3).

101. Zhang D, Qian J, Zhang P, Li H, Shen H, Li X, et al. Gasdermin D serves as a key executioner of pyroptosis in experimental cerebral ischemia and reperfusion model both in vivo and in vitro. J Neurosci Res. 2019;97(6):645–60.

102. Wald G. Molecular basis of visual excitation. Science. 1968;162(3850):230–9.

103. Kiser PD, Golczak M, Maeda A, Palczewski K. Key enzymes of the retinoid (visual) cycle in vertebrate retina. Biochim Biophys Acta. 2012;1821(1):137–51.

104. Janecke AR, Thompson DA, Utermann G, Becker C, Hubner CA, Schmid E, et al. Mutations in RDH12 encoding a photoreceptor cell retinol dehydrogenase cause childhood-onset severe retinal dystrophy. Nat Genet. 2004;36(8):850–4.

105. Gu SM, Thompson DA, Srikumari CR, Lorenz B, Finckh U, Nicoletti A, et al. Mutations in RPE65 cause autosomal recessive childhood-onset severe retinal dystrophy. Nat Genet. 1997;17(2):194–7.

106. Allikmets R, Singh N, Sun H, Shroyer NF, Hutchinson A, Chidambaram A, et al. A photoreceptor cell-specific ATP-binding transporter gene (ABCR) is mutated in recessive Stargardt macular dystrophy. Nat Genet. 1997;15(3):236–46.

107. Stefanovic Z, Mijailovic B, Arneric S, Stefanovic M, Mikuska M. [Immunofluorescent tests in chronic liver diseases]. Lijec Vjesn. 1975;97(1):11–5.

108. Ong DE, Davis JT, O’Day WT, Bok D. Synthesis and secretion of retinol-binding protein and transthyretin by cultured retinal pigment epithelium. Biochemistry. 1994;33(7):1835–42.

109. Soprano DR, Soprano KJ, Goodman DS. Retinol-binding protein messenger RNA levels in the liver and in extrahepatic tissues of the rat. J Lipid Res. 1986;27(2):166–71.

110. Kanai M, Raz A, Goodman DS. Retinol-binding protein: the transport protein for vitamin A in human plasma. J Clin Invest. 1968;47(9):2025–44.

111. Cukras C, Gaasterland T, Lee P, Gudiseva HV, Chavali VR, Pullakhandam R, et al. Exome analysis identified a novel mutation in the RBP4 gene in a consanguineous pedigree with retinal dystrophy and developmental abnormalities. PloS one. 2012;7(11):e50205.

112. Seeliger MW, Biesalski HK, Wissinger B, Gollnick H, Gielen S, Frank J, et al. Phenotype in retinol deficiency due to a hereditary defect in retinol binding protein synthesis. Investigative ophthalmology & visual science. 1999;40(1):3–11.

113. Liu L, Suzuki T, Shen J, Wakana S, Araki K, Yamamura KI, et al. Rescue of retinal morphology and function in a humanized mouse at the mouse retinol-binding protein locus. Lab Invest. 2017;97(4):395–408.

114. Quadro L, Blaner WS, Hamberger L, Van Gelder RN, Vogel S, Piantedosi R, et al. Muscle expression of human retinol-binding protein (RBP). Suppression of the visual defect of RBP knockout mice. J Biol Chem. 2002;277(33):30191–7.

115. Quadro L, Blaner WS, Salchow DJ, Vogel S, Piantedosi R, Gouras P, et al. Impaired retinal function and vitamin A availability in mice lacking retinol-binding protein. EMBO J. 1999;18(17):4633–44.

116. Summers JA, Harper AR, Feasley CL, Van-Der-Wel H, Byrum JN, Hermann M, et al. Identification of Apolipoprotein A-I as a Retinoic Acid-binding Protein in the Eye. J Biol Chem. 2016;291(36):18991–9005.

117. Welty FK. Hypobetalipoproteinemia and abetalipoproteinemia. Curr Opin Lipidol. 2014;25(3):161–8.

118. Piatigorsky J. Enigma of the abundant water-soluble cytoplasmic proteins of the cornea: the “refracton” hypothesis. Cornea. 2001;20(8):853–8.

119. Pappa A, Sophos NA, Vasiliou V. Corneal and stomach expression of aldehyde dehydrogenases: from fish to mammals. Chem Biol Interact. 2001;130-132(1-3):181–91.

120. Verhagen C, Hoekzema R, Verjans GM, Kijlstra A. Identification of bovine corneal protein 54 (BCP 54) as an aldehyde dehydrogenase. Experimental eye research. 1991;53(2):283–4.

121. Estey T, Chen Y, Carpenter JF, Vasiliou V. Structural and functional modifications of corneal crystallin ALDH3A1 by UVB light. PloS one. 2010;5(12):e15218.

122. Estey T, Piatigorsky J, Lassen N, Vasiliou V. ALDH3A1: a corneal crystallin with diverse functions. Experimental eye research. 2007;84(1):3–12.

123. Lassen N, Bateman JB, Estey T, Kuszak JR, Nees DW, Piatigorsky J, et al. Multiple and additive functions of ALDH3A1 and ALDH1A1: cataract phenotype and ocular oxidative damage in Aldh3a1(-/-)/Aldh1a1(-/-) knock-out mice. J Biol Chem. 2007;282(35):25668–76.

124. Kim SW, Lee J, Lee B, Rhim T. Proteomic analysis in pterygium; upregulated protein expression of ALDH3A1, PDIA3, and PRDX2. Mol Vis. 2014;20:1192-202.

125. Estey T, Cantore M, Weston PA, Carpenter JF, Petrash JM, Vasiliou V. Mechanisms involved in the protection of UV-induced protein inactivation by the corneal crystallin ALDH3A1. J Biol Chem. 2007;282(7):4382–92.

126. Uma L, Hariharan J, Sharma Y, Balasubramanian D. Corneal aldehyde dehydrogenase displays antioxidant properties. Experimental eye research. 1996;63(1):117–20.

127. Duester G. Alcohol dehydrogenase as a critical mediator of retinoic acid synthesis from vitamin A in the mouse embryo. J Nutr. 1998;128(2 Suppl):459S–62S.

128. Mootha VV, Kanoff JM, Shankardas J, Dimitrijevich S. Marked reduction of alcohol dehydrogenase in keratoconus corneal fibroblasts. Mol Vis. 2009;15:706-12.

129. Segre J. Complex redundancy to build a simple epidermal permeability barrier. Curr Opin Cell Biol. 2003;15(6):776–82.

130. Bernerd F, Asselineau D. Successive alteration and recovery of epidermal differentiation and morphogenesis after specific UVB-damages in skin reconstructed in vitro. Dev Biol. 1997;183(2):123–38.

131. Marshall D, Hardman MJ, Nield KM, Byrne C. Differentially expressed late constituents of the epidermal cornified envelope. Proc Natl Acad Sci U S A. 2001;98(23):13031–6.

132. Nakamura T, Nishida K, Dota A, Matsuki M, Yamanishi K, Kinoshita S. Elevated expression of transglutaminase 1 and keratinization-related proteins in conjunctiva in severe ocular surface disease. Investigative ophthalmology & visual science. 2001;42(3):549–56.

133. Castro-Munozledo F. Development of a spontaneous permanent cell line of rabbit corneal epithelial cells that undergoes sequential stages of differentiation in cell culture. J Cell Sci. 1994;107 (Pt 8):2343–51.

134. Tong L, Corrales RM, Chen Z, Villarreal AL, De Paiva CS, Beuerman R, et al. Expression and regulation of cornified envelope proteins in human corneal epithelium. Investigative ophthalmology & visual science. 2006;47(5):1938–46.

135. Turner HC, Budak MT, Akinci MA, Wolosin JM. Comparative analysis of human conjunctival and corneal epithelial gene expression with oligonucleotide microarrays. Investigative ophthalmology & visual science. 2007;48(5):2050–61.

136. Waseem A, Alam Y, Dogan B, White KN, Leigh IM, Waseem NH. Isolation, sequence and expression of the gene encoding human keratin 13. Gene. 1998;215(2):269–79.

137. Ramirez-Miranda A, Nakatsu MN, Zarei-Ghanavati S, Nguyen CV, Deng SX. Keratin 13 is a more specific marker of conjunctival epithelium than keratin 19. Mol Vis. 2011;17:1652-61.

138. Albers KM. Keratin biochemistry. Clin Dermatol. 1996;14(4):309–20.

139. Kim TH, Sung SE, Cheal Yoo J, Park JY, Yi GS, Heo JY, et al. Copine1 regulates neural stem cell functions during brain development. Biochem Biophys Res Commun. 2018;495(1):168–73.

140. Chen CK, Burns ME, He W, Wensel TG, Baylor DA, Simon MI. Slowed recovery of rod photoresponse in mice lacking the GTPase accelerating protein RGS9-1. Nature. 2000;403(6769):557–60.

141. GrandPre T, Nakamura F, Vartanian T, Strittmatter SM. Identification of the Nogo inhibitor of axon regeneration as a Reticulon protein. Nature. 2000;403(6768):439–44.

142. Oertle T, van der Haar ME, Bandtlow CE, Robeva A, Burfeind P, Buss A, et al. Nogo-A inhibits neurite outgrowth and cell spreading with three discrete regions. J Neurosci. 2003;23(13):5393–406.

143. Yang YS, Strittmatter SM. The reticulons: a family of proteins with diverse functions. Genome Biol. 2007;8(12):234.

144. Tagami S, Eguchi Y, Kinoshita M, Takeda M, Tsujimoto Y. A novel protein, RTN-XS, interacts with both Bcl-XL and Bcl-2 on endoplasmic reticulum and reduces their anti-apoptotic activity. Oncogene. 2000;19(50):5736–46.

145. Jerkovic L, Voegele AF, Chwatal S, Kronenberg F, Radcliffe CM, Wormald MR, et al. Afamin is a novel human vitamin E-binding glycoprotein characterization and in vitro expression. J Proteome Res. 2005;4(3):889–99.

146. Lichenstein HS, Lyons DE, Wurfel MM, Johnson DA, McGinley MD, Leidli JC, et al. Afamin is a new member of the albumin, alpha-fetoprotein, and vitamin D-binding protein gene family. J Biol Chem. 1994;269(27):18149–54.

147. Heiser M, Hutter-Paier B, Jerkovic L, Pfragner R, Windisch M, Becker-Andre M, et al. Vitamin E binding protein afamin protects neuronal cells in vitro. J Neural Transm Suppl. 2002(62):337–45.

148. Kratzer I, Bernhart E, Wintersperger A, Hammer A, Waltl S, Malle E, et al. Afamin is synthesized by cerebrovascular endothelial cells and mediates alpha-tocopherol transport across an in vitro model of the blood-brain barrier. J Neurochem. 2009;108(3):707–18.

149. Steele FR, Chader GJ, Johnson LV, Tombran-Tink J. Pigment epithelium-derived factor: neurotrophic activity and identification as a member of the serine protease inhibitor gene family. Proc Natl Acad Sci U S A. 1993;90(4):1526–30.

150. Dawson DW, Volpert OV, Gillis P, Crawford SE, Xu H, Benedict W, et al. Pigment epithelium-derived factor: a potent inhibitor of angiogenesis. Science. 1999;285(5425):245–8.

151. Ramirez-Castillejo C, Sanchez-Sanchez F, Andreu-Agullo C, Ferron SR, Aroca-Aguilar JD, Sanchez P, et al. Pigment epithelium-derived factor is a niche signal for neural stem cell renewal. Nat Neurosci. 2006;9(3):331–9.

152. Sugita Y, Becerra SP, Chader GJ, Schwartz JP. Pigment epithelium-derived factor (PEDF) has direct effects on the metabolism and proliferation of microglia and indirect effects on astrocytes. J Neurosci Res. 1997;49(6):710–8.

153. Taniwaki T, Hirashima N, Becerra SP, Chader GJ, Etcheberrigaray R, Schwartz JP. Pigment epithelium-derived factor protects cultured cerebellar granule cells against glutamate-induced neurotoxicity. J Neurochem. 1997;68(1):26–32.

154. Taniwaki T, Becerra SP, Chader GJ, Schwartz JP. Pigment epithelium-derived factor is a survival factor for cerebellar granule cells in culture. J Neurochem. 1995;64(6):2509–17.

155. Sun J, Wu, Zhong H, Guan D, Zhang H, Tan Q, et al. Presence of Zika Virus in Conjunctival Fluid. JAMA ophthalmology. 2016;134(11):1330–2.

156. Furtado JM, Esposito DL, Klein TM, Teixeira-Pinto T, da Fonseca BA. Uveitis Associated with Zika Virus Infection. N Engl J Med. 2016;375(4):394–6.

157. Slater L, Bartlett NW, Haas JJ, Zhu J, Message SD, Walton RP, et al. Co-ordinated role of TLR3, RIG-I and MDA5 in the innate response to rhinovirus in bronchial epithelium. PLoS pathogens. 2010;6(11):e1001178.

158. Gitlin L, Barchet W, Gilfillan S, Cella M, Beutler B, Flavell RA, et al. Essential role of mda-5 in type I IFN responses to polyriboinosinic:polyribocytidylic acid and encephalomyocarditis picornavirus. Proc Natl Acad Sci U S A. 2006;103(22):8459–64.

159. Loo YM, Gale M, Jr. Immune signaling by RIG-I-like receptors. Immunity. 2011;34(5):680–92.

160. Asgari S, Schlapbach LJ, Anchisi S, Hammer C, Bartha I, Junier T, et al. Severe viral respiratory infections in children with IFIH1 loss-of-function mutations. Proc Natl Acad Sci U S A. 2017;114(31):8342–7.

161. Walker CL, Little ME, Roby JA, Armistead B, Gale M, Jr., Rajagopal L, et al. Zika virus and the nonmicrocephalic fetus: why we should still worry. Am J Obstet Gynecol. 2019;220(1):45–56.

162. Aragao M, Holanda AC, Brainer-Lima AM, Petribu NCL, Castillo M, van der Linden V, et al. Nonmicrocephalic Infants with Congenital Zika Syndrome Suspected Only after Neuroimaging Evaluation Compared with Those with Microcephaly at Birth and Postnatally: How Large Is the Zika Virus “Iceberg”? AJNR Am J Neuroradiol. 2017;38(7):1427–34.

163. Linden VV, Linden HVJ, Leal MC, Rolim ELF, Linden AV, Aragao M, et al. Discordant clinical outcomes of congenital Zika virus infection in twin pregnancies. Arq Neuropsiquiatr. 2017;75(6):381–6.

164. Nguyen CTO, Hui F, Charng J, Velaedan S, van Koeverden AK, Lim JKH, et al. Retinal biomarkers provide “insight” into cortical pharmacology and disease. Pharmacol Ther. 2017;175:151-77.

165. van Wijngaarden P, Hadoux X, Alwan M, Keel S, Dirani M. Emerging ocular biomarkers of Alzheimer disease. Clin Exp Ophthalmol. 2017;45(1):54–61.

166. Dehghani C, Frost S, Jayasena R, Masters CL, Kanagasingam Y. Ocular Biomarkers of Alzheimer’s Disease: The Role of Anterior Eye and Potential Future Directions. Investigative ophthalmology & visual science. 2018;59(8):3554–63.

167. Zhou L, Beuerman RW. Tear analysis in ocular surface diseases. Prog Retin Eye Res. 2012;31(6):527–50.

168. Wei Y, Gadaria-Rathod N, Epstein S, Asbell P. Tear cytokine profile as a noninvasive biomarker of inflammation for ocular surface diseases: standard operating procedures. Investigative ophthalmology & visual science. 2013;54(13):8327–36.

169. Roy NS, Wei Y, Kuklinski E, Asbell PA. The Growing Need for Validated Biomarkers and Endpoints for Dry Eye Clinical Research. Investigative ophthalmology & visual science. 2017;58(6):BIO1–BIO19.

